# *C. elegans* PEZO-1 is a Mechanosensitive Ion Channel Involved in Food Sensation

**DOI:** 10.1101/2021.05.10.443141

**Authors:** Jonathan RM Millet, Luis O Romero, Jungsoo Lee, Briar Bell, Valeria Vásquez

## Abstract

PIEZO channels are force sensors essential for physiological processes, including baroreception and proprioception. The *Caenorhabditis elegans* genome encodes an ortholog gene of the *Piezo* family, *pezo-1*, which is expressed in several tissues, including the pharynx. This myogenic pump is an essential component of the *C. elegans* alimentary canal, whose contraction and relaxation are modulated by mechanical stimulation elicited by food content. Whether *pezo-1* encodes a mechanosensitive ion channel and contributes to pharyngeal function remain unknown. Here, we leverage genome editing, genetics, microfluidics, and electropharyngeogram recording to establish that *pezo-1* is expressed in the pharynx, including in a proprioceptive-like neuron, and regulates pharyngeal function. Knockout (KO) and gain-of-function (GOF) mutants reveal that *pezo-1* is involved in fine-tuning pharyngeal pumping frequency, sensing osmolarity, and food mechanical properties. Using pressure-clamp experiments in primary *C. elegans* embryo cultures, we determine that *pezo-1* KO cells do not display mechanosensitive currents, whereas cells expressing wild-type or GOF PEZO-1 exhibit mechanosensitivity. Moreover, infecting the *Spodoptera frugiperda* cell line with a baculovirus containing the G*-*isoform of *pezo-1* (among the longest isoforms) demonstrates that *pezo-1* encodes a mechanosensitive channel. Our findings reveal that *pezo-1* is a mechanosensitive ion channel that regulates food sensation in worms.

## INTRODUCTION

Mechanosensitive ion channels regulate several physiological processes ranging from osmotic balance in bacteria (Kung et al., 2010, Martinac et al., 1987), turgor control in plants (Hamilton et al., 2015), touch (Geffeney and Goodman, 2012, Yan et al., 2013, Ikeda et al., 2014, Maksimovic et al., 2014, Ranade et al., 2014, Woo et al., 2014, Chesler et al., 2016), pain (Murthy et al., 2018, Szczot et al., 2018), proprioception (Woo et al., 2015), hearing (Pan et al., 2018), lineage choice (Pathak et al., 2014), to blood pressure regulation in mammals (Retailleau et al., 2015, Wang et al., 2016, Rode et al., 2017, Zeng et al., 2018). These channels are ubiquitous, as they transduce mechanical stimuli into electrochemical signals in all kingdoms of life (Douguet and Honore, 2019, Geffeney and Goodman, 2012, Kung et al., 2010). In 2010, the PIEZO1 and PIEZO2 channels were identified as essential components of distinct, mechanically activated cation channels in mammalian cells (Coste et al., 2010). Since then, many physiological roles have been assigned to these two ion channels (Parpaite and Coste, 2017).

Mammalian PIEZO channels have been associated with several hereditary pathophysiologies (Alper, 2017). *Piezo1* gain-of-function (GOF) mutations display slow channel inactivation, leading to an increase in cation permeability and subsequent red blood cell dehydration (Albuisson et al., 2013, Bae et al., 2013, Ma et al., 2018, Zarychanski et al., 2012). For instance, the human *Piezo1* hereditary mutation R2456H, located in the pore domain, decreases inactivation, while substitution by Lys decreases inactivation even further (Bae et al., 2013). *Piezo1* global knockouts (KO) are embryonically lethal in mice (Li et al., 2014, Ranade et al., 2014) and cell-specific KOs result in animals with severe defects (Ma et al., 2018, Wu et al., 2017a). Intriguingly, both *Piezo2* KO and GOF mutations are associated with joint contractures, skeletal abnormalities and alterations in muscle tone (Chesler et al., 2016, Coste et al., 2013, Yamaguchi et al., 2019). GOF and loss-of-function (LOF) mutations are useful genetic tools for determining the contribution of PIEZO channels to mechanosensation in diverse physiological processes and in various animals.

The *C. elegans* genome encodes an ortholog of the PIEZO channel family, namely *pezo-1* (wormbase.org v. WS280). Recently, Bai and collaborators showed that *pezo-1* is expressed in several tissues, including the pharynx (Bai et al., 2020). The worm’s pharynx is a pumping organ that rhythmically couples muscle contraction and relaxation in a swallowing motion to pass food down to the animal’s intestine (Keane and Avery, 2003). This swallowing motion stems from a constant low-frequency pumping, maintained by pharyngeal muscles, and bursts of high-frequency pumping from a dedicated pharyngeal nervous system (Avery and Horvitz, 1989, Lee et al., 2017, Raizen et al., 1995, Trojanowski et al., 2016). In mammals, the swallowing reflex is initiated when pressure receptors in the pharynx walls are stimulated by food or liquids, yet the identity of the receptor(s) that directly evoke this mechanical response remain to be identified (Tsujimura et al., 2019). Interestingly, the *Drosophila melanogaster* PIEZO ortholog is a mechanosensitive ion channel (Kim et al., 2012) required for feeding, while also important for avoiding food over-consumption (Min et al., 2021, Wang et al., 2020). To date, whether *pezo-1* encodes for a mechanosensitive ion channel or regulates worm pharyngeal activity has yet to be determined.

Here, we found a strong and diverse expression of the *pezo-1* gene in pharyngeal tissues by imaging a *pezo-1::GFP* transgenic reporter strain. By leveraging genetic dissection, electrophysiological measurements, and behavior analyses, we also established that PEZO-1 is required for proper low frequency electrical activity and pumping behavior. Analyses of *pezo-1* KO and GOF mutants demonstrated that decreasing or increasing PEZO-1 function upregulates pharyngeal parameters. Likewise, mutants display distinct pharyngeal activities triggered by the neurotransmitter serotonin or with various buffer osmolarities. Using elongated bacteria as a food source, we demonstrated that *pezo-1* KO decreases pharyngeal pumping frequency, whereas a GOF mutant features increased frequency. Finally, electrophysiological recordings of *pezo-1* expressing cells from *C. elegans* embryo cultures and the *Spodoptera frugiperda* (Sf9) cell line demonstrate that *pezo-1* encodes a mechanosensitive ion channel. Altogether, our results reveal that PEZO-1 is a mechanosensitive ion channel involved in a novel biological function, regulating pharyngeal pumping and food sensation.

## MATERIALS AND METHODS

### Strains and Maintenance

Worms were propagated as previously described (Brenner, 1974). N2 (var. Bristol) is referred as wild type (WT) throughout the manuscript. The following strains were used: VVR3 *unc119(ed3)III;decEx1(pRedFlpHgr)(C10C5.1[20789]::S0001_pR6K_Amp_2xTY1ce_EGFP_FRT_ rpsl_neo_FRT_3xFlag)dFRT::unc-119-Na*t, COP1553 (KO: 6,616 bp deletion) *pezo-1 (knu508) IV*, COP1524 (GOF: R2373K) *pezo-1 (knu490) IV,* LX960 *lin-15B&lin-15A(n765) X*; *vsIs97 [tph-1p::DsRed2 + lin-15(+)],* DA572 *eat-4(ad572) III,* and DA1051 *avr-15(ad1051) V*. Transgenic strain VVR3 was obtained by microinjecting a fosmid construct (from the TransgeneOme Project) into a *unc-119(ed3)* strain (InVivo Biosystems). COP1553 and COP1524 were obtained using the CRISPR-Cas9 method (InVivo Biosystems). Transgenic worm VVR3, expressing GFP under the control of *Ppezo-1::GFP,* was crossed with *pezo-1* mutants COP1553 and COP1524 to obtain VVR69 and VVR70, respectively. LX960 was kindly provided by Dr. Kevin Collins (University of Miami).

### Genomic DNA PCR

PCR was performed with worm lysates of N2 and COP1553. Phusion High-Fidelity DNA Polymerase (ThermoScientific) was used to amplify *pezo-1* DNA fragments. Primer sequences are as follows: F1 5’-GACGACCAACTGCCTACAT-3’, F2 5’-TGCTGTAAATTAGCACTCGGGT-3’, and R1 5’-TGTACCGTAATCCAGAAACGCA-3’

### RNA isolation and RT PCR

Cultured worms were washed with M9 buffer (86 mM NaCl, 42 mM Na2HPO4, 22 mM KH2PO4, 1mM MgSO4) and collected into 15ml Falcon tubes. Trizol was added to pelleted worms and RNA was isolated using the freeze-cracking method as previously described (Van Gilst et al., 2005). Isolated total RNA was purified using RNAeasy kit (Qiagen). The SuperScript III One-Step RT-PCR system with Platinum Taq DNA Polymerase was used for RT-PCR, following the manufacturer’s protocol (Invitrogen). Primers were designed based on the *pezo-1* sequences comprising the deletion regions of the *knu508* allele of the COP1553 mutant: F3 5’-GCAACGTCACCAAGAAGAGCAG-3’ and R2 5’-GCATTCAATAGTCTCGTTGCTG-3’.

### Imaging

Worms were individually selected and dropped in 15 μL of M9 buffer, then paralyzed on a glass slide with 2% agarose pads containing 150 mM 2,3-butanedione monoxime (BDM). Brightfield and fluorescence imaging were performed on a Zeiss 710 Confocal microscope using a 20X or 40X objective. Images were processed using Fiji ImageJ (Schindelin et al., 2012) to enhance contrast and convert to an appropriate format.

### Worm synchronization

For all pharyngeal pumping assays, worms were synchronized by placing young adults onto fresh nematode growth media (NGM) plates seeded with OP50 (*E. coli* strain) and left to lay eggs for two hours at 20°C. Then, adults were removed, and the plates were incubated at 20°C for three days.

### Pharyngeal pumping

#### Serotonin profile

A serotonin aliquot (InVivo Biosystems) was diluted in M9 Buffer prior to experiments and discarded after three hours. 42 synchronized worms were picked and transferred to 200 μL of M9 Buffer supplemented with 2-, 5-, 10- or 20-mM serotonin and incubated at 20°C for 30 minutes before loading onto a microfluidic chip (SC40, The ScreenChip™ System, InVivo Biosystems).

#### Control E. coli assay

OP50 was grown in liquid LB medium under sterile conditions at 37°C and diluted to an optical density of 1.0. Bacterial cultures were stored at 4°C for up to one week.

#### Spaghetti-like E. coli assay

The day before the experiment, OP50 colonies were picked from a fresh LB plate and incubated in 2 mL of LB overnight. The following day, 0.5 mL of this culture was used to inoculate 1.5 mL of LB media and incubated until growth was exponential, which was verified by optical density (OD of 0.5). Then, cephalexin (Alfa Aesar) was added to a final concentration of 60 μg/ml and the culture was incubated for two hours. Spaghetti-like OP50 were verified under a microscope and washed three times using 2 mL of M9 buffer, followed by centrifugation at 400 *g* to gently pellet the elongated bacteria.

#### Pharyngeal recordings and analyses

Worms were loaded one-by-one into the microfluidic chip recording channel and left to adjust for one minute prior to recording. All recordings were two minutes long. Records were analyzed using NemAnalysis software (InVivo Biosystems) with the brute force algorithm turned off. Parameters were adjusted for each record to include the maximum number of clearly identifiable pharyngeal pumps. Results were exported from the software in sheet form and parameters were plotted and statistically analyzed using MATLAB R2019a (MathWorks).

### Development assay

Young adults were allowed to lay eggs on NGM plates seeded with control or spaghetti-like bacteria for two hours. Spaghetti-like bacteria were cultured as described above. Animals (10-20 worms) were removed from plates after two hours and the number of eggs laid was counted. After 3 days of incubation, animals that reached adulthood were counted in each trial and results were compared across four trials.

### Food ingestion assay

A drop of fresh culture containing control or spaghetti-like bacteria with 2 μM DiI dye (Sigma, CAS #41085-99-8) was placed on an NGM agar plate. Young adults were fed bacteria with DiI for 30 min. Next, worms were transferred onto OP50 seeded NGM without dye for 5 min (Vidal-Gadea et al., 2012). Finally, animals were placed on a thin-layered BDM-agarose plate for imaging under a Nikon SMZ18 stereomicroscope. Food occupation in the digestive tract was detected by fluorescence.

### Primary culture of *C. elegans* embryo cells

*C. elegans* embryonic cells were generated as previously described (Strange et al., 2007). Worms were grown on 10 cm enriched peptone plates with NA22 *E. coli*. NA22 bacteria grow in very thick layers that provide an abundant food source for large quantities of worms. Synchronized gravid hermaphrodites were bleached to release eggs and washed with sterile egg buffer (118 mM NaCl, 48 mM KCl, 2 mM CaCl_2_, 2 mM MgCl_2_, 25 mM HEPES, pH 7.3, 340 mOsm, adjusted with sucrose). The isolated eggs were separated from debris by centrifugation in a 30% sucrose solution. Chitinase (1 U/ml, Sigma) digestion was performed to remove eggshells. The embryo cells were dissociated by pipetting and filtered through a sterile 5μm Durapore filter (Millipore). The cells were plated on glass coverslips coated with peanut lectin solution (SIGMA; 0.5 mg/ml) and cultured in L15 media (Gibco) supplemented with 50 U/ml penicillin, 50 μg/ml streptomycin, and 10% fetal bovine serum (FBS, Invitrogen) for 72-96 hrs.

### Expression of *pezo-1* in Sf9 insect cells

To express *pezo-1* in Sf9 cells, we produced a recombinant baculovirus, according to the manufacturer’s instructions (Bac-to-Bac expression system, Invitrogen). To generate this baculovirus, we used a pFastBac construct (Epoch Life Science) containing an 8×histidine-maltose binding protein (MBP) tag and a synthesized *pezo-1* isoform G nucleotide sequence (one of the longest isoforms according to RNA sequencing, wormbase.org v. WS280). For expression of *pezo-1* R2373K, the construct contained an 8×histidine-MBP tag and a synthesized *pezo-1* isoform G with the R2373K point mutation. We infected Sf9 cells with either *pezo-1* baculovirus for 48 hours. Infected cells were plated on glass coverslips coated with a peanut lectin solution (SIGMA; 0.5 mg/ml) for patch-clamp experiments.

### Electrophysiology and mechanical stimulation

Primary cultured embryo cells labeled with *Ppezo-1::GFP* from strains VVR3, VVR69, or VVR70 were recorded in the cell-attached configuration of the patch-clamp technique. Control and infected Sf9 insect cells were recorded in the whole-cell or inside-out patch-clamp configurations. For on-cell recordings, the bath solution contained 140 mM KCl, 6 mM NaCl, 2 mM CaCl_2_, 1 mM MgCl_2_, 10 mM glucose, and 10 mM HEPES (pH 7.4; 340 mOsm, adjusted with sucrose). The pipette solution contained 140 mM NaCl, 6 mM KCl, 2 mM CaCl_2_, 1 mM MgCl_2_, 10 mM glucose, and 10 mM HEPES (pH 7.3; 330 mOsm, adjusted with sucrose). Cells were mechanically stimulated with negative pressure applied through the patch pipette using a High-Speed Pressure Clamp (ALA Scientific) controlled with a MultiClamp 700B amplifier through Clampex (Molecular Devices, LLC). Cell-attached patches were probed using a square-pulse protocol consisting of −10 mmHg incremental pressure steps, each lasting 1 s, in 10 s intervals. Cells with giga-seals that did not withstand at least six consecutive steps of mechanical stimulation were excluded from analyses. I_steady_ was defined as the maximal current in the steady state. Deactivation was compared by determining the percentage of I_steady_ remianing 100 ms after removal of the mechanical stimuli.

For whole-cell recordings, the bath solution contained 140 mM NaCl, 6 mM KCl, 2 mM CaCl_2_, 1 mM MgCl_2_, 10 mM glucose, and 10 mM HEPES (pH 7.4). The pipette solution contained 140 mM CsCl, 5 mM EGTA, 1 mM CaCl_2_, 1 mM MgCl_2_, and 10 mM HEPES (pH 7.2). For indentation assays, Sf9 cells were mechanically stimulated with a heat-polished blunt glass pipette (3–4 μm) driven by a piezo servo controller (E625, Physik Instrumente). The blunt pipette was mounted on a micromanipulator at an ~45° angle and positioned 3–4 μm above the cells without indenting them. Displacement measurements were obtained with a square-pulse protocol consisting of 1 μm incremental indentation steps, each lasting 200 ms, with a 2-ms ramp in 10 s intervals. Recordings with leak currents >200 pA, with access resistance >10 MΩ, and cells with giga-seals that did not withstand at least five consecutive steps of mechanical stimulation were excluded from analyses. For inside-out recordings, symmetrical conditions were established with a solution containing 140 mM CsCl, 5 mM EGTA, 1 mM CaCl_2_, 1 mM MgCl_2_, and 10 mM HEPES (pH 7.2), and mechanical stimulation was performed identically to on-cell recordings. Pipettes were made from borosilicate glass (Sutter Instruments) and were fire-polished, before use, until a resistance between 3 and 4 MΩ was reached. Currents were recorded at a constant voltage (−60 mV, unless otherwise noticed), sampled at 20 kHz, and low pass filtered at 2 kHz using a MultiClamp 700B amplifier and Clampex (Molecular Devices, LLC). Leak currents before mechanical stimulation were subtracted offline from the current traces.

Data and fits were plotted using OriginPro (from OriginLab). Sigmoidal fit was done with the following Boltzmann equation:

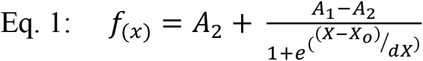

where A_2_ = final value, A_1_ = initial value; X_o_ = center, and *dX* = time constant.

The time constant of inactivation τ was obtained by fitting a single exponential function, Equation (2), between the peak value of the current and the end of the stimulus:

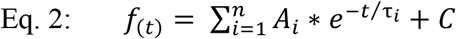

where A = amplitude, τ = time constant, and the constant y-offset *C* for each component *i*.

### Data and Statistical analyses

Data and statistical analyses were performed using DataGraph 4.6.1, MATLAB R2019a (MathWorks), and GraphPad Instat 3 software. Statistical methods and sample numbers are detailed in the corresponding figure legends. No technical replicates were included in the analyses.

### Data availability

Data supporting the findings of this manuscript are available from the corresponding author upon reasonable request. The source data underlying Figures and Supplementary Figures are provided as a Source Data file, DOI: 10.6084/m9.figshare.16992058.

## RESULTS

### *pezo-1* is expressed in a wide variety of cells in the worm pharynx

To determine the expression of *pezo-1* in *C. elegans*, we used a fluorescent translational reporter made by the TransgeneOme Project (Hasse et al., 2016). This fosmid construct contains *pezo-1* native cis-regulatory elements, including introns, up to exon 17 and 3’ UTR sequences linked in-frame to the green fluorescent protein (GFP; Fig. 1 A). The position of the GFP with respect to the remainder of the gene creates an unnatural truncated version of the PEZO-1 protein. Hence, it likely expresses a non-functional protein that excludes 16 exons, which contain most of the *pezo-1* sequence (including the pore domain). GFP signals are present in all developmental stages and multiple cells (Fig. S1 A-B). Furthermore, it does not appear to be mosaic as similar expression patterns were observed in at least three independent transgenic lines. We imaged *pezo-1::GFP* worms at various focal planes to identify the different cells expressing GFP based on their morphological features (i.e., cell-body position, neurites extension and position along the body, and branching) (Fig. 1 B–G). The strongest GFP signals that we identified come from the pharyngeal gland cells (Fig. 1 B, bright and fluorescence fields). These cells are composed of five cell bodies (two ventral g1s, one dorsal g1 and two ventral g2s) located inside the pharynx terminal bulb and three anterior cytoplasmic projections: two short projections that are superposed, ending in the metacorpus, and a long projection reaching the end of the pm3 muscle. These cells are proposed to be involved in digestion (Albertson and Thomson, 1976, Ohmachi et al., 1999), lubrication of the pharynx (Smit et al., 2008), generation and molting of the cuticle (Hoflich et al., 2004, Singh, 1978), and resistance to pathogenic bacteria (Hoflich et al., 2004). Additionally, we visualized *pezo-1::GFP* in a series of cells surrounding the muscle of the corpus and the isthmus (Fig. 1 C) (Albertson and Thomson, 1976). We also recognized the arcade cells as putative *pezo-1* expressing cells, according to their morphology and location (Fig. 1 C and E). Arcade cells and the pharyngeal epithelium form the buccal cavity and connect the digestive tract to the outside (Altun, 2013). We also observed many other anterior cells labeled with GFP; however, we cannot currently confirm if they represent neurons and/or amphids.

**Figure 1.**
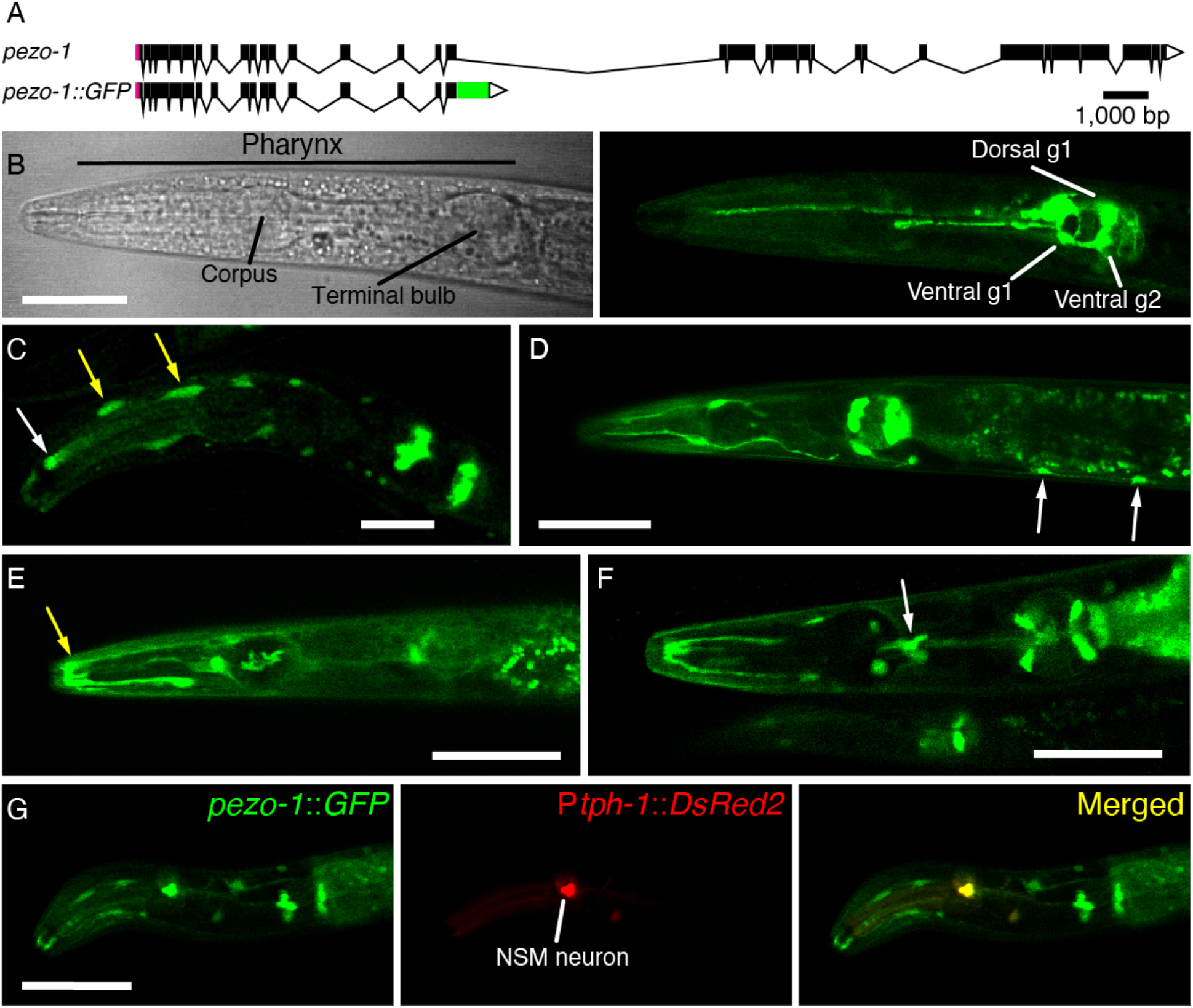
*pezo-1* is strongly expressed in *C. elegans* pharynx. (**A**) *pezo-1* gene diagram according to wormbase.org, v. WS280 made with Exon-Intron Graphic Maker (wormweb.org). Magenta rectangles and white triangles denote the 5’ and 3’ untranslated regions (UTR), respectively. Black rectangles denote exons and black lines denote introns. Green rectangle denotes the GFP sequence inserted after exon 17. (**B**) Brightfield (left) and fluorescence (right) micrographs of the anterior end of a young adult *pezo-1*::*GFP* hermaphrodite highlighting pharynx structures and the GFP reporter expression in gland cells. Scale bar represents 50 μm. (**C**) Micrograph of the anterior end of a young adult *pezo-1*::*GFP* hermaphrodite expressing GFP in different cells. Arrows highlight pm3 muscle (white) and arcade cells (yellow), according to their morphology and location. Scale bar represents 20 μm. (**D-E**) Micrograph of the anterior end of a young adult *pezo-1*::*GFP* hermaphrodite expressing GFP in different cells. White and yellow arrows highlight VNC neurons and posterior arcade syncytium, respectively, according to their morphology and location. Scale bar represents 50 μm. (**F**) Micrograph of the anterior end of a young adult *pezo-1*::*GFP* hermaphrodite expressing GFP in the pharyngeal sieve (white arrow), according to its morphology and location. Scale bar represents 100 μm. (**G**) Co-localization between *tph-1*::DsRed2 and *pezo-1*::*GFP* reporter in the NSM neuron. Scale bar represents 50 μm. Micrographs are representative of at least 20 independent preparations.

By crossing *pezo-1::GFP* with a *tph-1*::DsRed marker carrying strain, we were able to identify *pezo-1* expression in the pharyngeal NSM_L/R_ secretory, motor, and sensory neurons (Fig. 1 G). Importantly, these serotoninergic neurons have been proposed to sense food in the lumen of the pharynx through their proprioceptive-like endings and trigger feeding-related behaviors (i.e., increased pharyngeal pumping, decreased locomotion, and increased egg laying) (Albertson and Thomson, 1976, Avery et al., 1993). In addition to the pharyngeal cells, we observed expression of *pezo-1* in the ventral nerve cord (VNC) neurons, according to their morphology and location (Fig. 1 D), striated muscles (Fig. S1 C), coelomocytes (Fig. S1 D-E), spermatheca (Fig. S1 B and 1E), vulval muscles (Fig. S1 F), and various male neurons, including the ray neurons (Fig. S1 G). Importantly, the expression pattern reported by our *pezo-1* fosmid construct in NSM neurons matches with what is reported in the *C. elegans* Neuronal Gene Expression Map & Network (CeNGEN; (Taylor et al., 2021). The strong and varied *pezo-1* expression in the pharynx, along with the function of the cells expressing it, led us to investigate the potential contribution of PEZO-1 to pharyngeal function.

### Serotonin stimulation reveals varying pharyngeal pump parameters

To analyze the contribution of *pezo-1* to pharyngeal pumping in *C. elegans*, we used the ScreenChip™ system (InVivo Biosystems) that allows measuring electropharyngeogram recording (EPG; Fig. 2 A) (Raizen and Avery, 1994) by loading single, live worms into a microfluidic chip. Fig. 2 A-B summarizes the pharynx anatomy, electrical properties measured during an EPG, and the neurons involved in pharyngeal function. For instance, the excitation event (E spike) precedes the pharyngeal contraction and is modulated by the MC pacemaker neuron (Fig. 2 B, top), whereas the repolarization event (R spike) leads to pharyngeal relaxation and correlates with the activity of the inhibitory M3 motor neurons (Fig. 2 B, middle). Every 3-4 pumps, there is relaxation of the terminal bulb (isthmus peristalsis), which is modulated by the M4 motor neuron (Fig. 2 B, bottom) (Avery and Horvitz, 1989). The main EPG events are regulated by the pharyngeal proprioceptive neuron, NSM. Importantly, the proprioceptive NSM neurons and the I3 interneuron express *pezo-1* according to our data (Fig. 1 G) and CeNGEN (Taylor et al., 2021), respectively.

**Figure 2.**
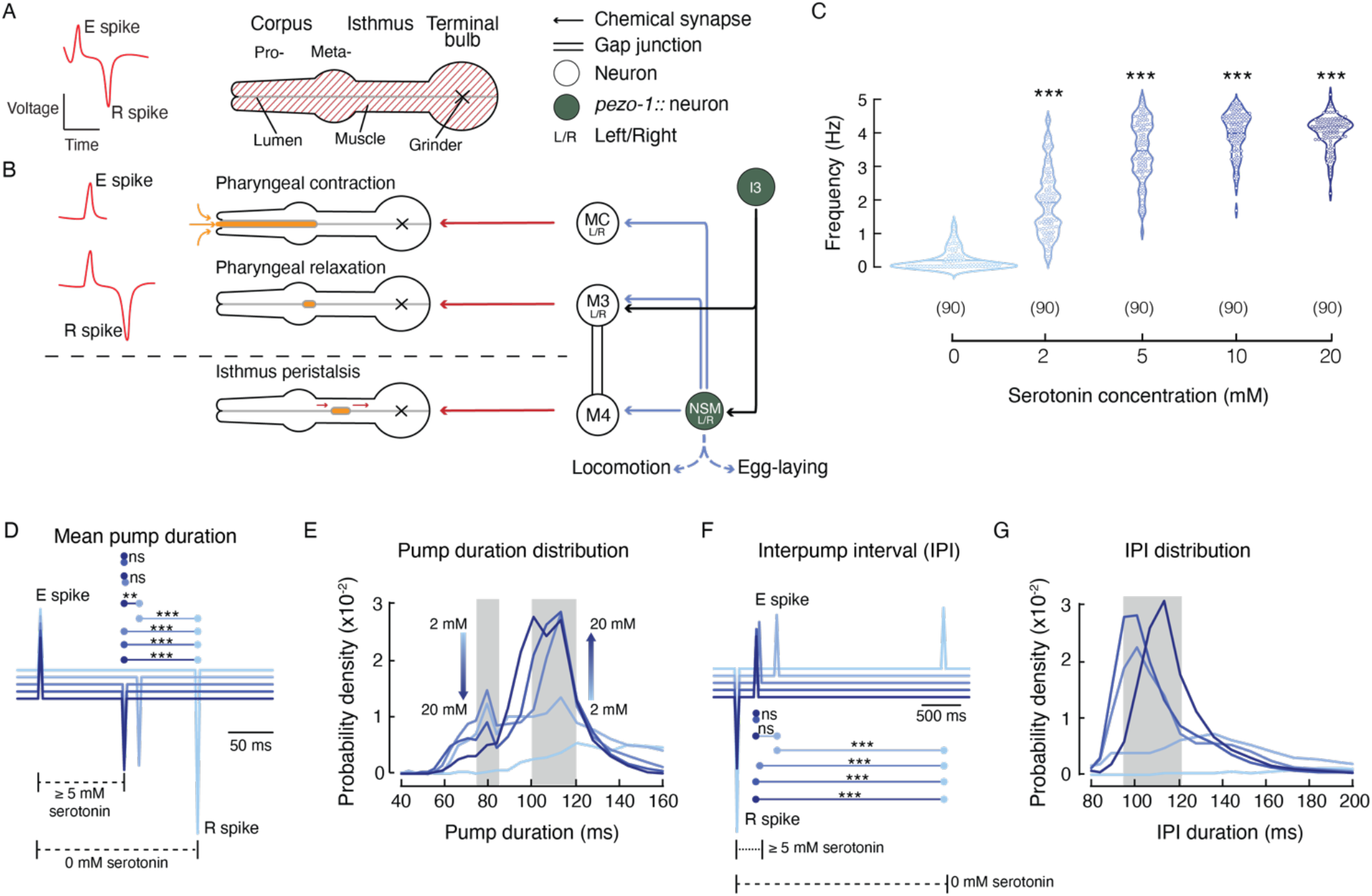
Serotonin triggers pharyngeal pumping in a concentration-dependent manner in WT (N2) worms. (**A**) Left, illustration of an electropharyngeogram (EPG) recording depicting the contraction-relaxation of the corpus, referred to as pharyngeal pump. Right, illustration representing the *C. elegans* pharyngeal muscle. (**B**) Representation of the main pharyngeal electrical coupling in *C. elegans*. Blue arrows represent serotoninergic transmission, red arrows represent neuromuscular junctions, double lines represent gap junctions, filled circles represent the neurons expressing *pezo-1* (NSM and I3 according to our data and CeNGEN (Taylor et al., 2021), respectively). (**C**) Pharyngeal pumping frequencies, depicted as violin plots with the means shown as horizontal bars, for WT (N2) at different serotonin concentrations. Kruskal-Wallis and Dunn’s multiple comparisons tests (0 mM vs. 2, 5, 10, 20 mM. n is denoted above the *x*-axis. (**D**) Ensemble averages of EPG traces for different serotonin concentrations showing pump durations (E to R spikes). One-way ANOVA and Tukey-Kramer multiple comparisons test. n is 90 worms per condition (**E**) Serotonin concentration effect on pump duration. A kernel probability distribution was fit to the data. n is 90 worms per condition. (**F**) Ensemble averages of EPG traces for different serotonin concentrations showing interpump interval (R to E spikes). One-way ANOVA and Tukey-Kramer multiple comparisons test. n is 90 worms per condition (**G**) Serotonin concentration effect on interpump interval. A kernel probability distribution was fit to the data. n is 90 worms per condition. Asterisks indicate values significantly different (***p < 0.001 and **p < 0.01) and ns indicates not significantly different.

Analysis of the EPG records allows for the determination of various pharyngeal pumping parameters, including frequency, duration, and the time interval that separates two pumping events (hereafter referred to as the interpump interval). We used serotonin to increase pharyngeal activity since, in the absence of food or serotonin, pumping events are infrequent. Serotonin mimics food stimulation by activating the MC_L/R_ and M3_L/R_ neurons (Niacaris and Avery, 2003). First, we established a serotonin dose-response profile of the WT (N2) strain pharyngeal pumping parameters (Fig. 2 C–G). Serotonin increases pharyngeal pumping frequency in a dose-dependent manner, with concentrations above 5 mM increasing the likelihood of reaching 5 Hz (Fig. 2 C). We averaged the EPG recordings at each serotonin concentration and found a clear difference in pump duration between 0- and 5-mM. Concentrations equal to or higher than 5 mM evoke similar pump durations (~100 ms; Fig. 2 D). Interestingly, analysis of the pump duration distribution profile under serotonin stimulation revealed that pharyngeal pump duration fits into two categories: fast (~80 ms) and slow (100-120 ms; Fig. 2 E; gray rectangles).

We observed that the fast and slow categories displayed an inverse relationship with respect to serotonin concentration (Fig. 2 E; arrows). Unlike pump duration, we observed only a single category for interpump interval, around 95-120 ms for serotonin concentrations of 5- to 20-mM (Fig. 2 F–G). Interestingly, we did not observe interpump intervals faster than 90 ms, regardless of the serotonin concentration. The interpump interval results support the idea that there is a minimum refractory period between two pumps. This set of analyses allowed us to establish a suitable model for evaluating the role of *pezo-1* function *in vivo*.

### PEZO-1 modulates pump duration and interpump interval

To determine whether *pezo-1* has a functional role in pharyngeal pumping, we engineered LOF and GOF mutants. A putative LOF mutant was obtained by deleting 6,616 bp from the *pezo-1* locus (hereafter referred to as *pezo-1* KO; Fig. 3 A, top and Fig. S2 A-C). This CRISPR-KO transgenesis deleted 638 amino acid residues from PEZO-1 that, according to the cryo-EM structure of the PIEZO1 mouse ortholog (Ge et al., 2015, Guo and MacKinnon, 2017, Saotome et al., 2018, Zhao et al., 2018), eliminates 12 transmembrane segments, seven extra- and intra-cellular loops, and the beam helix that runs parallel to the plasma membrane (Fig. S2 D). Previous works demonstrated that the substitution of R2456H (located at the pore helix) in the ortholog human *Piezo1* gene increases cation permeability (GOF) and causes hemolytic anemia (Albuisson et al., 2013, Bae et al., 2013, Zarychanski et al., 2012). Moreover, a conservative substitution of Lys for Arg at position 2456 in the human PIEZO1 channel exhibits a pronounced slowed inactivation when compared to the WT or R2456H channels (Bae et al., 2013). Hence, we engineered a putative GOF mutant strain, obtained by substituting the conserved Arg 2373 with Lys (hereafter referred to as *pezo-1* R2373K or GOF; Fig. 3 A, bottom). Parenthetically, the R2373K numbering position is based on isoform G – one of the longest isoforms according to RNA sequencing (wormbase.org v. WS280). We also included two mutants known to alter pharyngeal function, *eat-4(ad572)* and *avr-15(ad1051)*, in our analysis. EAT-4 is a glutamate-sodium symporter involved in postsynaptic glutamate reuptake. *eat-4(ad572)* affects the neurotransmission efficiency of all glutamatergic pharyngeal neurons (I2L/R, I5, M3_L/R_, M4, MI, NSM_L/R_) (Lee et al., 1999). AVR-15 is a glutamate-gated chloride channel expressed in the pm4 and pm5 pharyngeal muscles (both synapsed by M3_L/R_) and is involved in relaxation of the pharynx. Its mutant allele *ad1051* lengthens pump duration by delaying relaxation of the pharynx in a similar fashion to laser ablation of M3_L/R_ neurons (Dent et al., 1997). With these strains, we sought to determine if altering PEZO-1 function would affect the worm’s pharyngeal phenotype.

**Figure 3.**
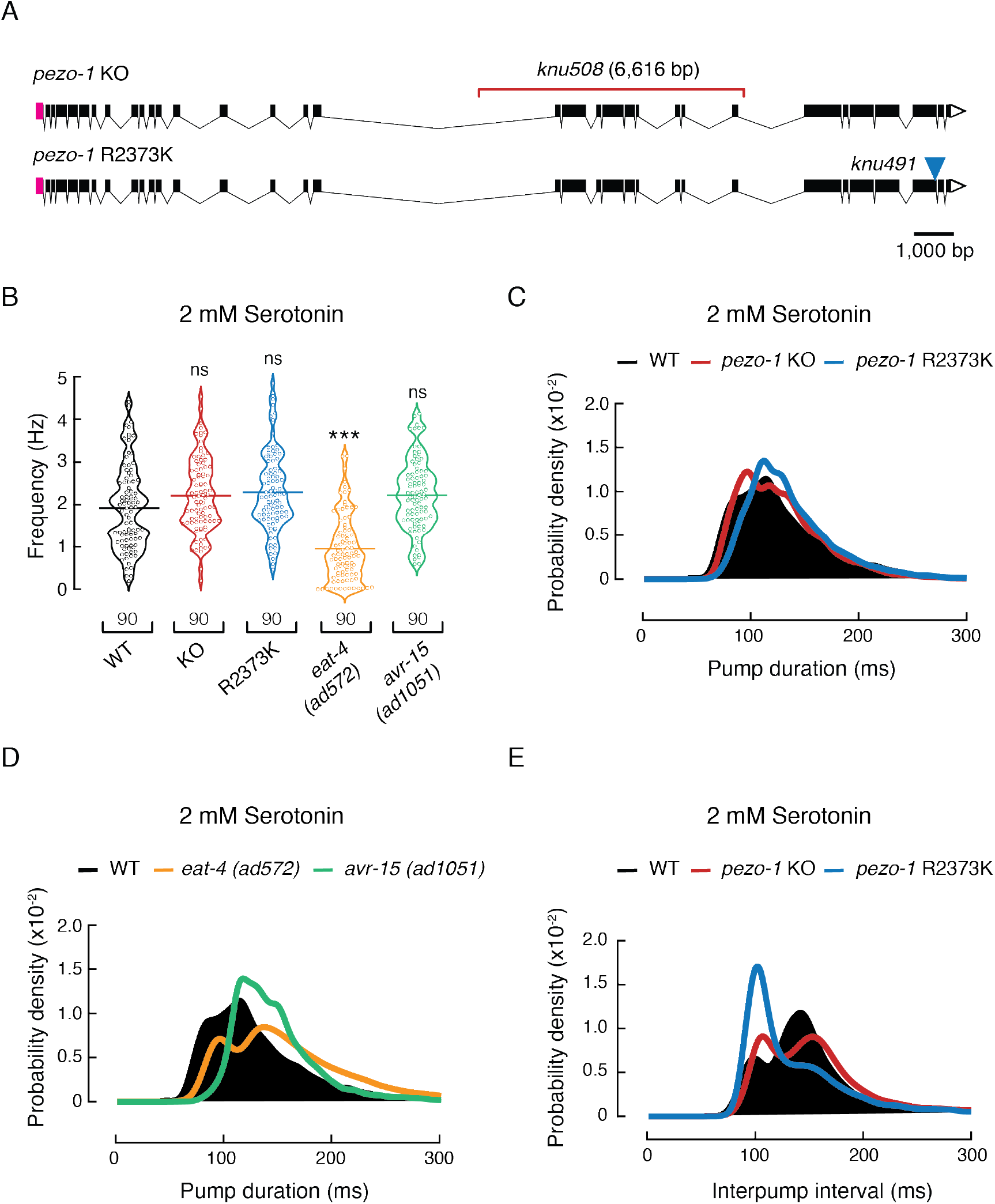
*pezo-1* modulates pharyngeal pumping properties at 2 mM serotonin. (**A**) *pezo-1* gene diagram according to wormbase.org v. WS280 made with Exon-Intron Graphic Maker (wormweb.org). Magenta rectangles and white triangles denote the 5’ and 3’ untranslated regions (UTR), respectively. Black rectangles denote exons and black lines denote introns. The red bracket denotes the *knu508* allele (a 6,616bp deletion) of the *pezo-1* KO strain and the blue triangle denotes the single point mutation (allele *knu491*) of the *pezo-1* R2373K strain. (**B**) Pharyngeal pumping frequencies depicted as violin plots with the means shown as horizontal bars for WT (N2), *pezo-1* KO, *pezo-1* R2373K, *eat-4 (ad572)*, and *avr-15 (ad1051)* strains. n is denoted above the *x*-axis. Kruskal-Wallis and Dunn’s multiple comparisons tests. Asterisks indicate values significantly different (***p < 0.001) and ns indicates not significant. (**C**) Kernel density plot of WT, *pezo-1* KO, and *pezo-1* R2373K strain pump durations. n is 90 worms per condition. (**D**) Kernel density plot of WT, *eat-4 (ad572)*, and *avr-15 (ad1051)* strain pump durations. n is 90 worms per condition. (**E**) Kernel density plot of WT, *pezo-1* KO, and *pezo-1* R2373K strain interpump intervals. n is 90 worms per condition.

At a 2 mM concentration of exogenous serotonin (to elicit pharyngeal activity), both *pezo-1* KO and R2373K mutants displayed higher pumping frequencies than WT, albeit not statistically significant (WT 1.92 ± 0.11 Hz, KO 2.21± 0.09 Hz, GOF 2.29 ± 0.1, 2.22 ± 0.09 Hz, mean ± SEM; Fig. 3 B), similar to *avr-15(ad1051)* (2.22 ± 0.09 Hz, mean ± SEM; Fig. 3 B). On the other hand, the *eat-4(ad572)* mutant displayed lower pumping frequency at this serotonin concentration. To further assess the altered pharyngeal function *pezo-1* mutants, we analyzed the pump duration distributions from the EPG records. *pezo-1* KO distribution is similar to the WT (Fig. 3 C, red *vs.* black), whereas the R2373K mutant profile is reminiscent of *avr-15(ad1051)*, as both mutant strains displayed significantly narrower distributions around 100 ms pump events (Fig. 3 C–D, blue and green *vs.* black), when compared to the WT (significance was determined by a Z-test). Moreover, the R2373K mutant lacked fast pump events, between 50 to 80 ms (Fig. 3 C), similar to the WT features observed at high serotonin concentrations (≥ 5 mM, Fig. 2 E), and the *eat-4(ad572)* and the *avr-15(ad1051)* mutants at a 2 mM serotonin concentration (Fig. 3 D). Analysis of the distribution of interpump intervals revealed that *pezo-1* KO and R2373K mutants, although different, both spend less time resting between pumps (95-120 ms) than the WT (≈ 140 ms) (Fig. 3 E). This enhancement in function resembles the WT activity measured at 5- to 20-mM serotonin concentrations (Fig. 2 F–G) and could account for the slight increase in frequency shown in Fig. 3 B. The close resemblance between the pharyngeal pumping parameters of PEZO-1 GOF and the *avr-15(ad1051)* mutant suggests a potential link between PEZO-1 and pharyngeal relaxation.

### PEZO-1 determines pharyngeal pumping in response to osmolarity

Mechanical stimuli come in many forms, including stretching, bending, and osmotic forces (Cox et al., 2019). To further understand the functional role of *pezo-1*, we evaluated pharyngeal pumping parameters after challenging worm strains with varying osmolarities (in the absence of serotonin or food). The worm’s pharynx draws in liquid and suspended bacteria from the environment, then expels the liquid but traps bacteria (Avery and You, 2012). We adjusted a standard laboratory solution used for worm experiments (M9 buffer, 320 mOsm) to varying osmolarities (150, 260, and 320 mOsm). Parenthetically, M9 buffer elicits acute withdrawal behavior in the absence of food, and another buffer (M13) with lower osmolarity (*~* 280 mOsm) is commonly used to study molecules that elicit acute avoidance behavior (Caires et al., 2021, Caires et al., 2017, Geron et al., 2018, Jang and Bargmann, 2013, Hart, 2006). Because we measured the pharynx function in the absence of food for this set of experiments, we refer to the M9 buffer as a “high osmolarity solution”. Low osmolarity solutions would be equivalent to swallowing liquid containing few solutes (150 mOsm), whereas high osmolarities would represent a “gulp” of liquid with a large amount of solutes (320 mOsm). Of note, higher osmolarities were associated with smaller mean pumping frequencies for WT worms (Fig. 4 A). Our results indicate that a larger number of solutes in solution corresponds to an increased retention time in the pharynx before moving to the intestine of the WT worms. Notably, at 320 mOsm, both *pezo-1* KO and GOF mutants displayed a significantly higher pumping frequency than WT worms (Fig. 4 A). In contrast, we did not measure significant differences between WT worms and the *pezo-1* mutants at 260 or 150 mOsm. Akin to human Piezo2 KO or GOF mutations causing joint contractures (Chesler et al., 2016, Coste et al., 2013, McMillin et al., 2014), we demonstrated that lack of or enhanced PEZO-1 function modulated pharyngeal pumping frequencies similarly (at high osmolarities). Next, we further examined the EPG parameters at high osmolarity (320 mOsm). Analysis of the distribution of pump durations and length of the mean interpump intervals revealed that both *pezo-1* mutants had more frequent fast pumps (80-120 ms, Fig. 4 B) and the KO spent less time resting between pumps, compared to the WT (Fig. 4 C). Interestingly, high osmolarity (320 mOsm) revealed resemblances between PEZO-1 GOF and *avr-15(ad1051) vs.* PEZO-1 KO and *eat-4(ad572)* pharyngeal pumping parameters (frequency and duration, Fig. 4 D–E). Altogether, our results suggest that PEZO-1 is required for fine tuning pharyngeal function in response to osmolarity changes.

**Figure 4.**
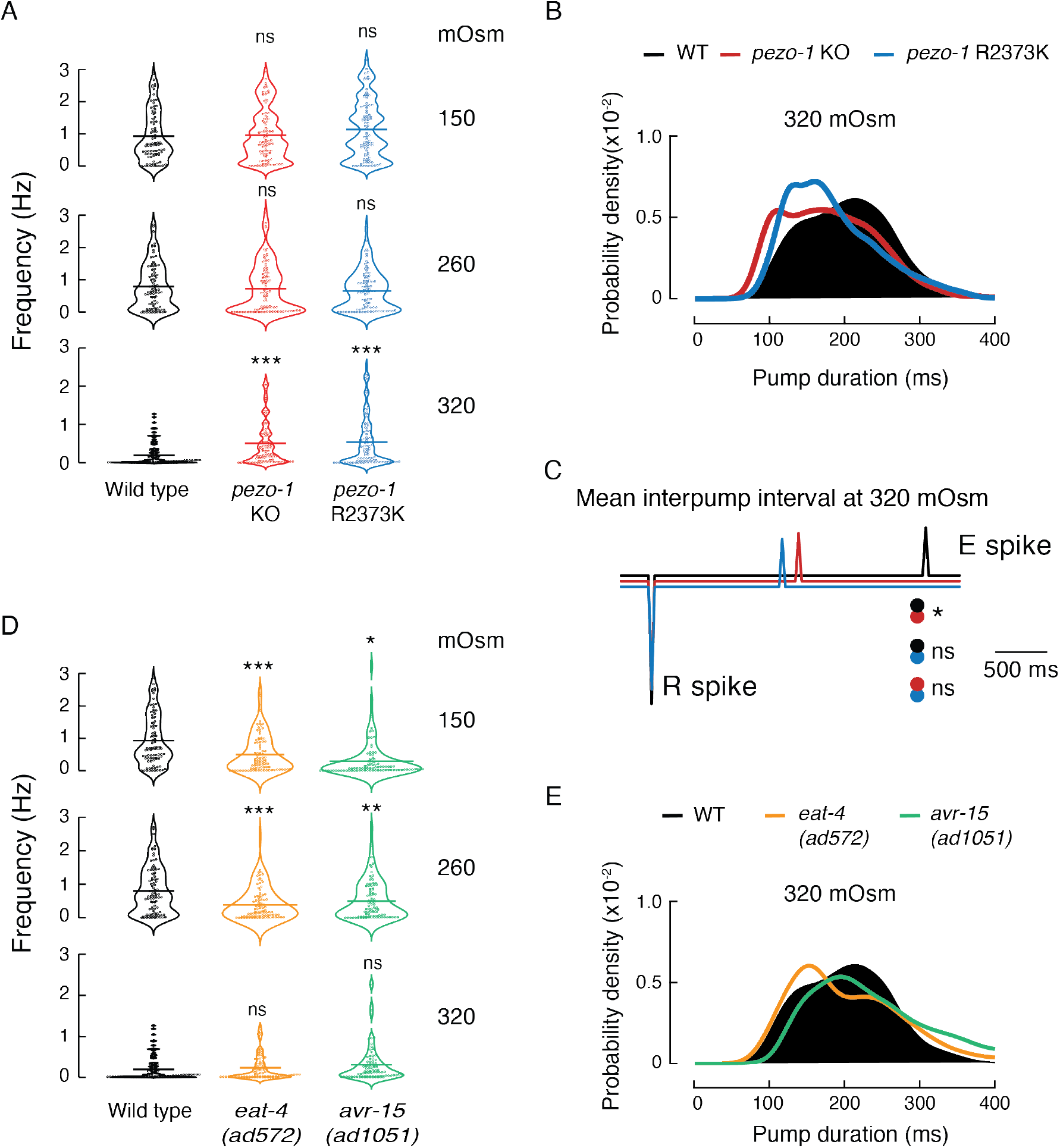
Osmolarity changes modify pharyngeal pumping in WT and *pezo-1* strains, in the absence of serotonin. (**A**) Pharyngeal pumping frequencies depicted as violin plots with the means shown as horizontal bars for WT (N2), *pezo-1* KO, and *pezo-1* R2373K strains at 150, 260, and 320 mOsm. n is 90 worms per condition. Kruskal-Wallis and Dunn’s multiple comparisons tests. (**B**) Kernel density plot of WT, *pezo-1* KO, and *pezo-1* R2373K strain pump durations. n is 90 worms per condition at 320 mOsm. (**C**) Ensemble averages of EPG traces for WT (N2), *pezo-1* KO, and *pezo-1* R2373K strains showing interpump intervals (R to E spikes) at 320 mOsm. One-way ANOVA and Tukey-Kramer multiple comparisons test. n is 90 worms per condition. (**D**) Pharyngeal pumping frequencies depicted as violin plots with the means shown as horizontal bars for WT (N2), *eat-4 (ad572)*, and *avr-15 (ad1051)* strains at 150, 260, and 320 mOsm. n is 90 worms per condition. Kruskal-Wallis and Dunn’s multiple comparisons tests. (**E**) Kernel density plot of WT, *eat-4 (ad572)*, and *avr-15 (ad1051)* strain pump durations at 320 mOsm. n is 90 worms per condition. Asterisks indicate values significantly different (***p < 0.001, **p < 0.01, and *p < 0.05) and ns indicates not significant.

### PEZO-1 function is involved in food sensation

To determine the impact that PEZO-1 function has on food intake, we recorded pharyngeal pumping of WT and *pezo-1* strains in response to varying food stimuli. It has been hypothesized that food quality and feeding preferences displayed by worms is linked to bacterial size (Shtonda and Avery, 2006). To this end, we measured worm pharyngeal pumping while feeding them with conventional food used in the laboratory for maintenance (*Escherichia coli* strain OP50). We found that feeding WT worms *E. coli* elicited lower pumping frequencies than 2 mM serotonin (Fig. S3; *E. coli* 1.36 ± 0.09 Hz and serotonin 1.92 ± 0.11 Hz, mean ± SEM), whereas *pezo-1* mutants displayed similar pumping frequencies with exogenous serotonin or *E. coli* (Fig. S3). Future experiments are needed to understand why exogenous serotonin stimulation and feeding result in varying *pezo-1* influence on pharyngeal function. Additionally, we varied the dimensions of OP50 using the antibiotic cephalexin, an antibiotic that prevents the separation of budding bacteria, which generates long spaghetti-like filaments of bacterium, as observed under a microscope and elsewhere (Hou et al., 2020, Martinac et al., 1987) (Fig. S4 A). Specifically, cephalexin yields bacterium whose contour length that is five to ten-times longer and 1.5 times stiffer (i.e., Young’s modulus and the bacterial spring constant) than untreated bacteria, as determined by fluorescence imaging and atomic force microscopy (Hou et al., 2020). Hence, feeding worms with these two physically different types of food could help to elucidate the physiological role of PEZO-1 in detecting food with varying mechanical properties (i.e., small and soft *vs.* large and rigid food). A similar method was previously described using the antibiotic aztreonam and was shown to affect pharyngeal pumping (Ben Arous et al., 2009, Gruninger et al., 2008). WT and *pezo-1* mutants are able to ingest spaghetti-like bacteria and reached adulthood in three days, similar to worms fed with control bacteria (Fig. S4 B-C). Notably, feeding worms with control or spaghetti-like bacteria revealed distinctive pharyngeal traits between the *pezo-1* mutants and the WT worms (Fig. S4 D). When fed with control *E. coli*, both *pezo-1* mutants (KO and GOF) have higher mean frequencies, shorter mean pump durations, narrower pump duration distributions, and faster mean interpump intervals than the WT worms (Fig. 5 A–E). Conversely, feeding worms with spaghetti-like *E. coli* elicits opposing effects on the pharyngeal pumping parameters of the *pezo-1* mutants. For instance, feeding with spaghetti-like *E. coli* decreases *pezo-1* KO mean frequency, while the mean pump duration and distribution remain similar to WT worms (Fig. 5 A–C). Furthermore, this modified diet significantly increases the mean interpump interval of the KO in comparison to the GOF mutant (Fig. 5 D). Unlike the KO and WT strains, the R2372K *pezo-1* mutant displays high frequency, shorter pumps (mean and distributions; Fig. 5 A–C) and reduced mean interpump interval durations (mean and distributions; Fig. 5 D–E). Altogether, our results indicate that PEZO-1 regulates the pharynx response to the physical parameters of food, such as the length and stiffness of ingested bacteria.

**Figure 5.**
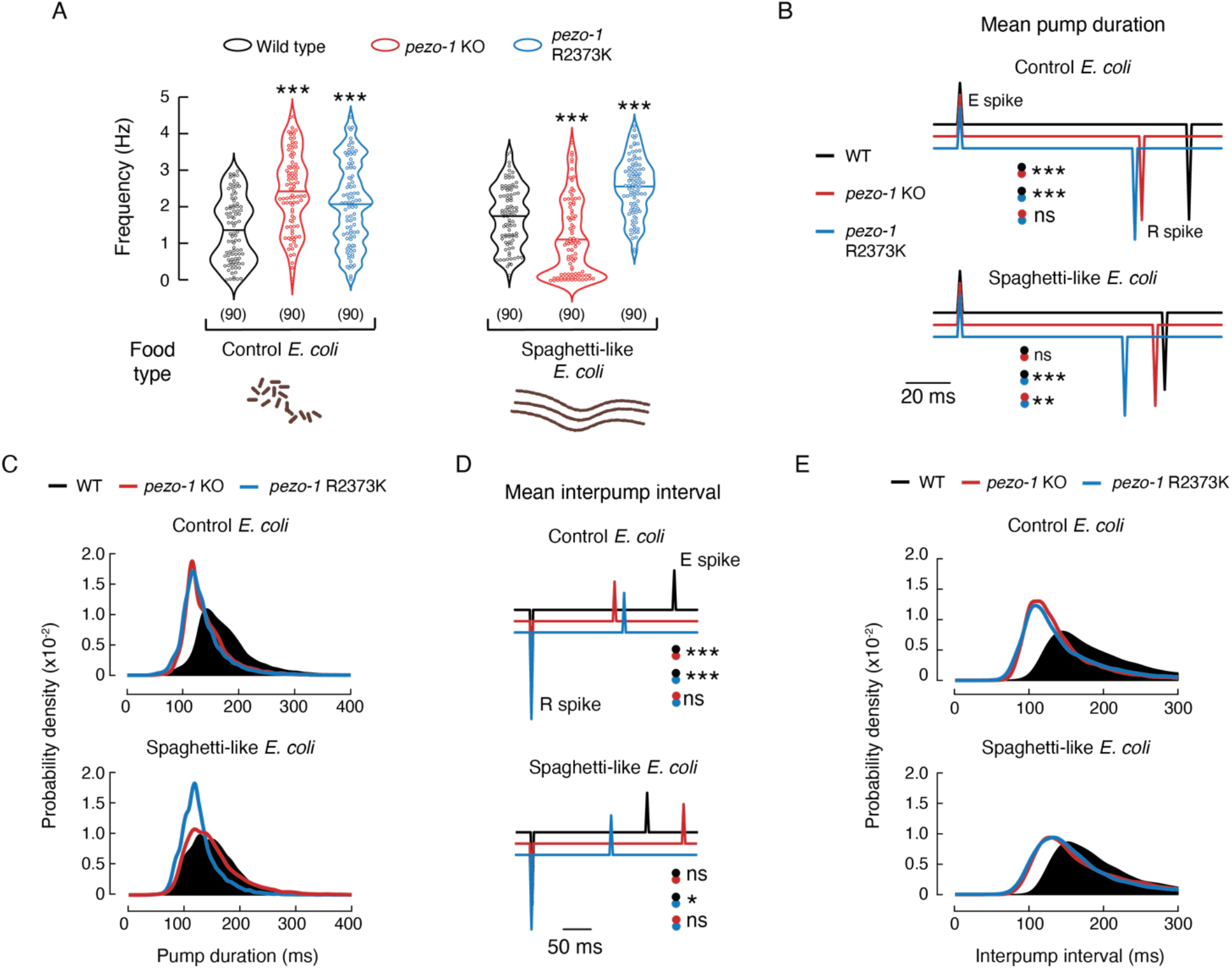
PEZO-1 alters pharyngeal pumping when exposed to control or spaghetti-like bacteria. (**A**) Pharyngeal pumping frequencies depicted as violin plots with the means shown as horizontal bars, for WT (N2), *pezo-1* KO, and *pezo-1* R2373K strains when fed with control or cephalexin-treated *E. coli* (spaghetti-like bacteria). Bacteria cartoon depictions were created with BioRender.com. n is denoted above the *x*-axis. Kruskal-Wallis and Dunn’s multiple comparisons tests. (**B**) Ensemble averages of EPG traces for WT (N2), *pezo-1* KO, and *pezo-1* R2373K strains showing pump durations (E to R spikes), when fed with control or cephalexin-treated bacteria (spaghetti-like *E. coli*). One-way ANOVA and Tukey-Kramer multiple comparisons test. n is 90 worms per condition. (**C**) Kernel density plot of WT, *pezo-1* KO, and *pezo-1* R2373K strains pump durations when fed with control or cephalexin-treated bacteria. n is 90 worms per condition. (**D)** Ensemble averages of EPG traces for WT (N2), *pezo-1* KO, and *pezo-1* R2373K strains showing interpump interval (R to E spikes), when fed with control or cephalexin-treated bacteria. One-way ANOVA and Tukey-Kramer multiple comparisons test. n is 90 worms per condition. (**E**) Kernel density plot of WT, *pezo-1* KO, and *pezo-1* R2373K strain interpump interval when fed with control or cephalexin-treated bacteria. n is 90 worms per condition. Asterisks indicate values significantly different (***p < 0.001, **p < 0.01, and *p < 0.05) and ns indicates not significantly different.

### *pezo-1* encodes a mechanosensitive ion channel

The PEZO-1 protein sequence shares 60-70% similarity with mammalian PIEZO channel orthologs. However, whether PEZO-1 responds to mechanical stimuli has not yet been established. To address this major question, we cultured *C. elegans* cells from three different strains endogenously expressing *pezo-1* WT, KO, or the R2373K GOF mutation in the background of the VVR3 strain that expresses a non-functional *pezo-1::GFP* reporter (Fig. 1 A). *pezo-1* WT, KO, and GOF strains express similar levels of GFP (Fig. S5). Embryonic *pezo-1::GFP* cells were patch-clamped using the cell-attached configuration, with application of constant negative pressure (−70 mmHg) at different voltages (Fig. 6 A–C). The normalized steady-state current (I/Imax) *vs.* voltage relationship is characterized by a reversal potential of + 9.06 mV (Fig. 6 D), indicating that PEZO-1 mediates a slight cation selective conductance, like the mouse and *Drosophila* orthologs (Coste, 2012). Importantly, most of the *pezo-1* WT cells displayed multiple-channel opening events upon mechanical stimulation. Nevertheless, from the few traces that carry pressure-dependent single-channel opening events, we were able to determine that PEZO-1 displays an outward slope conductance of 46.4 pS and an inward slope conductance of 34.8 pS (Fig. 6 E and Fig. S6). These current magnitudes are similar to the conductance of human PIEZO1 (37.1 pS) reported by others (Gottlieb et al., 2012).

**Figure 6.**
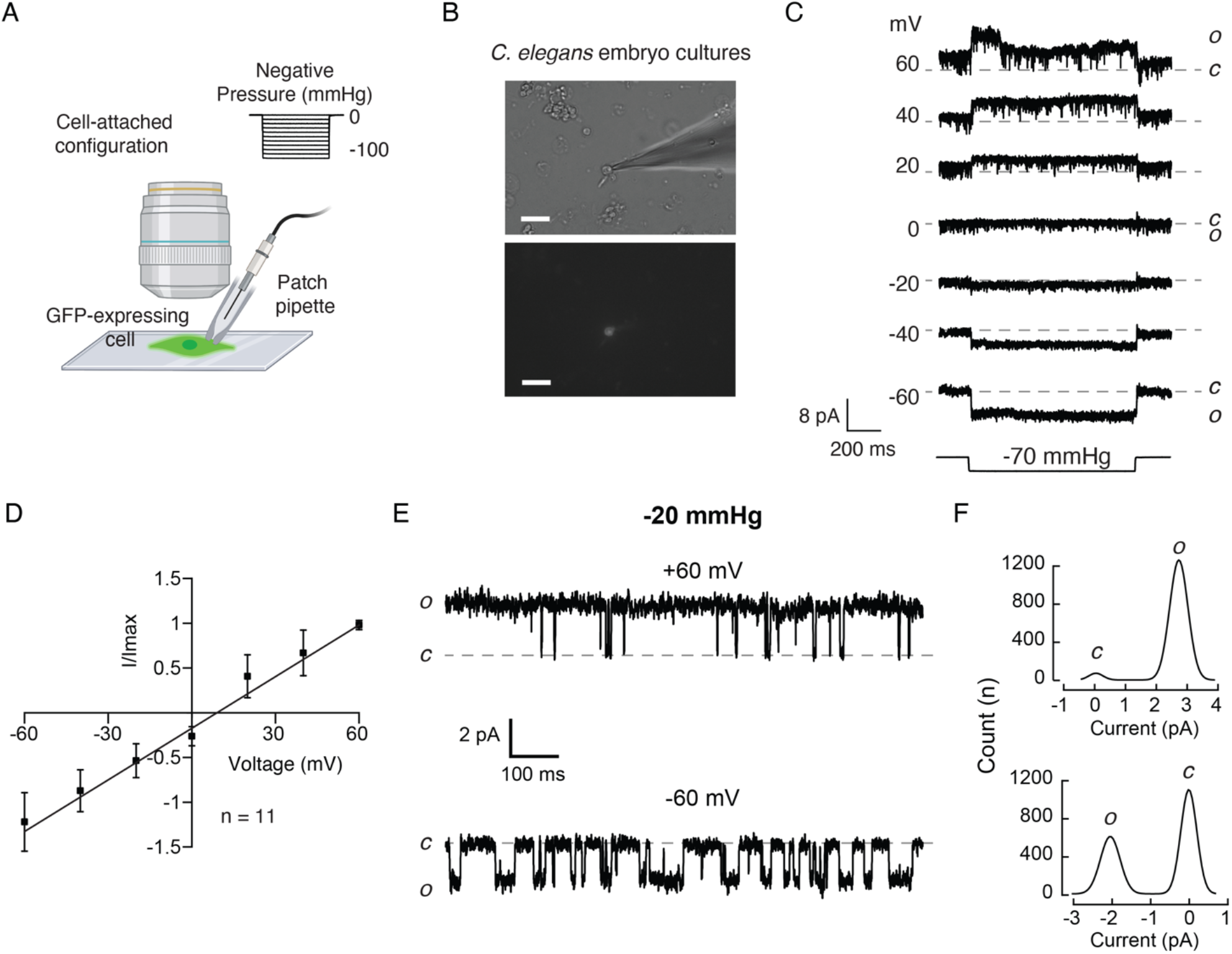
Cells expressing PEZO-1 display mechanosensitive channel currents. (**A**) Schematic representation of the mechanical stimulation protocol applied to *pezo-1*::*GFP* expressing cells recorded in the cell-attached configuration. Created with BioRender.com. (**B**) Representative micrographs (from at least 10 independent preparations) of a *C. elegans* primary embryonic culture from *pezo-1::GFP* expressing strains. Brightfield (top) and fluorescence (bottom) micrographs of a *pezo-1::GFP* expressing cell when patch-clamped in the on-cell configuration. Scale bar represents 20 μm. (**C**) Representative cell-attached patch-clamp recordings of mechanically-activated currents of WT cells expressing *pezo-1::GFP*. Channel openings were elicited by application of a −70 mmHg-square pulse (bottom) at constant voltages ranging from −60 to +60 mV. Dashed gray line represents background currents. Closed and open states are labeled *c* and *o*, respectively. **(D)** Normalized current-voltage relationship recorded at constant pressure in the cell-attached configuration. The reversal potential is 9.06 mV. Each square represents the mean ± SD. **(E)** Representative single-channel trace recordings of WT cells expressing *pezo-1::GFP* in the cell-attached configuration. Channel openings were elicited by −20 mmHg of negative pressure at constant voltages (+60 or −60 mV). Closed and open states are labeled *c* and *o*, respectively. **(F)** All-point amplitude histograms of pressure-evoked single-channel currents from recordings shown in E.

Mechanical stimulation of embryonic *pezo-1::GFP* cells expressing WT and GOF PEZO-1, but not the KO, yielded several channel openings upon increasing negative pressure (Fig. 7 A and 7C). As would be expected with increased activity, it is more difficult to identify single channel opening events in the traces coming from the GOF, as compared to the WT (Fig. 7 B and Fig. S7). Cells expressing WT PEZO-1 displayed mechano-dependent currents with a half pressure activation (P1/2) corresponding to −59.1 ± 4.3 mmHg (mean ± SEM; Fig. 7 A and 7 D). Alternatively, PEZO-1 R2373K displayed mechano-dependent currents with a significantly lower P1/2 than the WT channel (−39.2 ± 2.2 mmHg, mean ± SEM; Fig. 7 A and 7 D), indicating that the GOF mutant requires less mechanical stimulation to open. Since we could not reach saturating stimulus, these P1/2 values might be inaccurate; hence, future experiments are needed to unequivocally determine the differences in sensitivity between the WT and GOF. Notably, the R2373K mutation introduced a latency for activation that was not detected in the WT (Fig. 7 A, blue traces and 7 E). The decrease in mechanical threshold, along with the slowed activation, were previously reported for the equivalent human PIEZO1 R2456K mutation in mammalian cell lines (Albuisson et al., 2013, Bae et al., 2013, Romero et al., 2019, Zarychanski et al., 2012). Future experiments are needed to understand the origin of these differences in activation.

**Figure 7.**
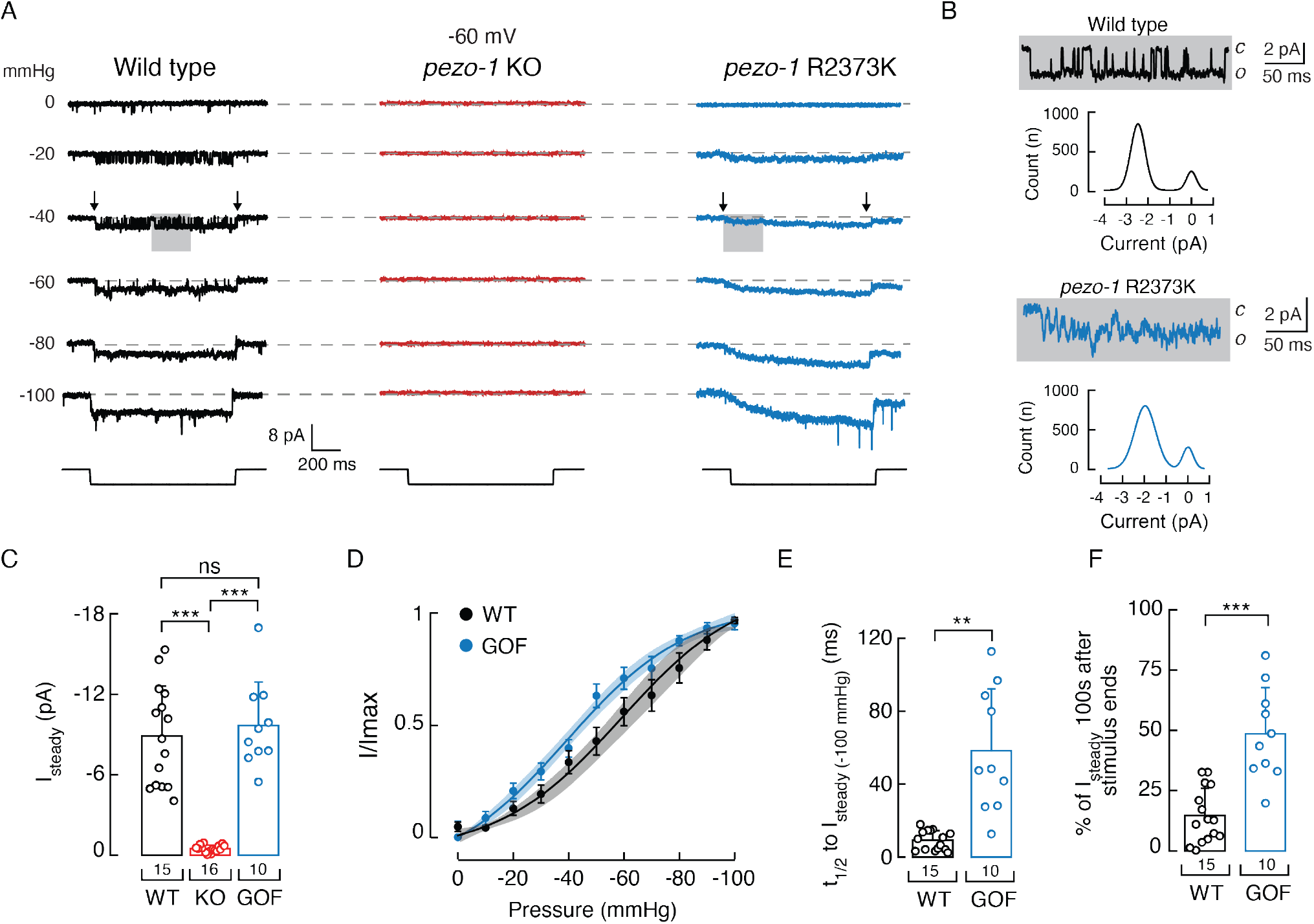
*pezo-1* GOF mutant decreases the mechanical threshold and slows down deactivation. (**A**) Representative cell-attached patch-clamp recordings of mechanically-activated currents from WT, *p*ezo-1 KO, and *pezo-1* R2373K cells expressing *pezo-1::GFP*. Channel openings (downward) were elicited by negative pressure (left) square pulses (bottom) at a constant voltage of −60 mV. Dashed gray line represents background currents. Gray rectangles highlight records magnified on panel B. Black arrows mark the recording sections (see filtered records on Supplementary Figure 7) used to generate the all-point amplitude histograms shown on panel B. (**B**) Gray rectangles show higher magnifications of indicated pressure-evoked responses shown in panel A. Traces are accompanied by all-point amplitude histograms generated from the corresponding records highlighted by arrows in panel A. (**C**) Bar graph displaying steady state currents elicited by −100 mmHg of negative pressure of WT, *p*ezo-1 KO, and *pezo-1* R2373K cells expressing *pezo-1::GFP*. Bars are mean ± SD. n is denoted above the *x*-axis. Kruskall-Wallis and Dunn’s multiple comparisons tests. (**D**) Pressure-response profiles for *pezo-1* WT and R2373K currents. Normalized currents (I/Imax) elicited by negative pressure of mechanically activated currents of WT and *pezo-1* R2373K cells expressing *pezo-1::GFP*. A Boltzmann function, Eq. (1), was fitted to the data. The shadows encompassing the curves indicate the 95% confidence bands for the fit. Circles are mean ± SD. n for WT and *pezo-1* R2373K are 15 and 10, respectively. (**E**) Bar graph displaying the time it takes to reach half of the steady state currents, elicited by −100 mmHg of pressure, of WT and *pezo-1* R2373K cells expressing *pezo-1::GFP*. Bars are all mean ± SD. n is denoted above the *x*-axis. Unpaired *t*-test with Welch’s correction. (**F**) Bar graph displaying the percentage of steady state currents remaining 100 ms after the removal of the mechanical stimulus, from WT and *pezo-1* R2373K cells expressing *pezo-1::GFP*. Bars are mean ± SD. n is denoted above the *x*-axis. Unpaired *t*-test. Asterisks (∗∗∗p < 0.001 and ∗∗p < 0.01) indicate values significantly different and ns indicates not significant.

Unlike *pezo-1* WT, approximately 50% of the mechanocurrents elicited from the *pezo-1* R2373K expressing cells remained active after removal of the mechanical stimulus (Fig. 7A, blue traces, 7F, and Fig. S7). This slow deactivation is also reminiscent of the human PIEZO1 R2456K GOF phenotype previously characterized by Bae and collaborators (Bae et al., 2013). Overall, our results suggest that PEZO-1 is a mechanosensitive ion channel and that a conservative mutation in the pore domain elicits similar activation and deactivation changes to its human counterpart.

One caveat of our PEZO-1 electrophysiological characterization in *C. elegans* cultured cells is that we cannot identify (at this point) which type of cells we are measuring from. We are only confident that those cells express *pezo-1* since they are labeled with the non-functional *pezo-1::GFP* reporter. These patch-clamp experiments were blind to genotype, yet we consistently found that *pezo-1::GFP* labeled cells coming from KO worms have negligible or no currents under the same voltage and pressure regimes used for *pezo-1 WT* and GOF cell cultures (Fig. 7 C). Hence, to further validate that the *pezo-1* gene encodes for a mechanosensitive ion channel, we heterologously expressed one of the longest isoforms of *pezo-1* (isoform G; wormbase.org v. WS280) in Sf9 cells and measured its function in the whole-cell patch-clamp configuration while stimulating with a piezo-electrically driven glass probe (Fig. 8 A). Similar to mammalian PIEZO channels, PEZO-1 mediates indentation-activated currents (Fig. 8 B). Uninfected Sf9 cells do not display mechanosensitive channel currents (Fig. 8 B and Fig. S8). Importantly PEZO-1 displayed the properties described for mammalian PIEZOs in other cell types (Coste et al., 2010, Wu et al., 2017b), including voltage-dependent inactivation (Fig. 8 C and 8 F) and non-selective cation currents, as determined by the reversal potential (−1.15 mV; Fig. 8 G). Our results demonstrate that expressing *pezo-1* in a naïve system was sufficient to confer mechanosensitivity to Sf9 cells.

**Figure 8.**
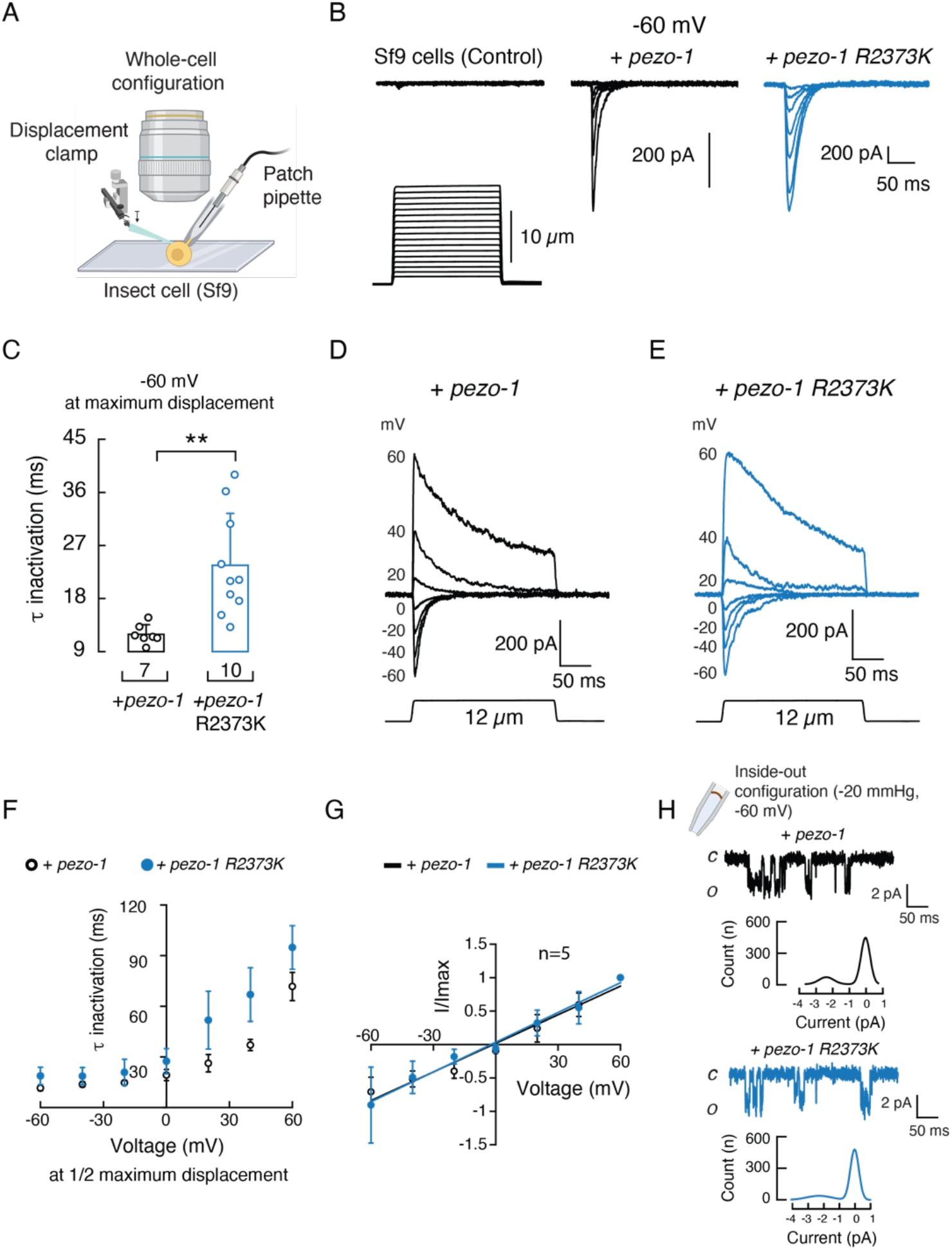
Sf9 cells infected with *pezo-1* WT- or *R2373K*-containing baculovirus display mechanosensitive currents. (**A**) Schematic representation of the mechanical stimulation protocol applied to Sf9 cells infected with baculovirus containing *pezo-1* (WT or R2373K constructs) recorded in the wholecell configuration. Created with BioRender.com. (**B**) Representative whole-cell patch-clamp recordings (at −60 mV) of currents elicited by mechanical stimulation of Sf9 cells, control or expressing *pezo-1* WT or R2373K. **(C)** Time constants of inactivation elicited by maximum displacement (-60 mV) of Sf9 cells expressing *pezo-1* WT or R2373K. Bars are mean ± SD. Two-tailed unpaired *t*-test with Welch correction. (**D**) Representative whole-cell patch-clamp recordings (top) at constant voltages (ranging from -60 to +60 mV) currents elicited by mechanical stimulation (bottom) of Sf9 cells expressing *pezo-1* WT. (**E**) Representative whole-cell patch-clamp recordings (top) at constant voltages (ranging from -60 to +60 mV) of currents elicited by mechanical stimulation (bottom) of Sf9 cells expressing *pezo-1* R2373K. (**F**) Time constants of inactivation elicited by ½ maximum displacement at constant voltages ranging from -60 to +60 mV, of Sf9 cells expressing *pezo-1* WT (n = 5) or R2373K (n = 5). Each circle represents the mean ± SD. (**G**) Normalized current-voltage relationship, recorded in the whole-cell configuration at a constant displacement of 12 μm. The reversal potential is -1.15 mV and -2.23 mV for *pezo-1* WT and R2373K, respectively. Each circle represents the mean ± SD. **(H)** Representative single-channel trace recordings and all-point amplitude histograms of pressure-evoked currents from Sf9 cells expressing *pezo-1* WT or R2373K, recorded in the inside-out configuration. Channel openings were elicited by -20 mmHg of negative pressure at -60 mV. Closed and open states are labeled *c* and *o*, respectively.

We also evaluated the PEZO-1 R2373K mutation in Sf9 cells and found that, like the human equivalent (Bae et al., 2013), the GOF elicits larger mechanically-evoked currents that inactivate more slowly than the WT channel (Fig. 8 B-F and Fig. S8). WT and GOF PEZO-1 channels are characterized by similar reversal potentials and unitary conductance, as determined by whole-cell and in inside-out patches, respectively (Fig. 8 G-H and Fig. S9). As previously reported with human PIEZO1 (Bae et al., 2013), we found that PEZO-1 (WT and GOF) features unique characteristics when activated with two different types of mechanical stimuli (negative pressure or displacement). For instance, with negative pressure, WT and mutant channels display steady-state currents, but only the GOF mutant exhibited a pronounced latency for activation and slowed deactivation. With displacement stimulation, it is possible to determine that PEZO-1 GOF inactivates more slowly than the WT. Overall, our results indicate that PIEZO orthologs are functionally conserved.

## DISCUSSION

In 2010, Coste and collaborators reported that the *C. elegans* genome contained a single *Piezo* gene, *pezo-1* (Coste et al., 2010). However, the functional role of *pezo-1* remained elusive, even a decade after its discovery. Here, we showed that PEZO-1 is a mechanosensitive channel with a novel functional role in the worm pharynx by combining fluorescent reporters, genome editing, electropharyngeogram recordings, behavior, and patch-clamp measurements. We found that *pezo-1* is highly expressed in gland cells and the proprioceptive-like NSM neuron, among many other tissues. In addition to its expression, several lines of evidence suggested that PEZO-1 modulated several discrete but reliable features of pharyngeal function. Lack- or augmentation of PEZO-1 function decreased interpump intervals when worms were challenged with 2 mM serotonin, while it also increased pumping frequency in high osmolarity conditions or feeding with control bacteria. In the absence of functional PEZO-1, worms reduced pharyngeal function (i.e., low frequency and long pump intervals) when fed with spaghetti-like bacteria. Finally, we demonstrated that the *pezo-*1 gene (WT or GOF) encodes a mechanosensitive ion channel by measuring its native function in *C. elegans* cells or with overexpression in insect cells. Altogether, our results established that PEZO-1 is important for pharyngeal function, regulation, and food sensation.

*C. elegans* feeding relies on the ability of its pharynx to contract and relax. The pharynx is a tube of electrically coupled muscle cells that continuously pump throughout the worm’s life (Mango, 2007). Several ion channels have been identified to be crucial for the pharyngeal muscle action potential, including acetylcholine receptors, T- and L-type Ca^2+^ channels, glycine receptors, and K^+^ channels (Avery and You, 2012). Although the pharyngeal muscle is capable of pumping (albeit at low frequencies) without nervous system input, higher pumping frequencies are controlled by pharyngeal motor neurons, namely MC_L/R_ and M3_L/R_ (Avery and You, 2012). Nevertheless, the role of the nervous system in the control of rhythmic pharyngeal pumping is not completely understood. It is known, however, that the pharynx responds to a variety of neuromodulators (Avery and Horvitz, 1989). We found that *pezo-1* is expressed in proprioceptive/mechanosensory NSM_L/R_ neurons (which are important for the pharyngeal nervous system). Moreover, Taylor and colleagues reported expression of *pezo-1* in the pharyngeal I3 interneuron (Taylor et al., 2021). Unlike NSM_L/R_ and M3_L/R_, the function of I3 has not been established (Avery, 1993, Avery and Thomas, 1997). Our results suggest that PEZO-1 is not essential for pharyngeal muscles, but instead fine-tunes the role of the nervous system in controlling pharynx function. This is reminiscent of the novel role of mammalian PIEZO1 and PIEZO2 in mediating neuronal sensing of blood pressure and the baroreceptor reflex (Zeng et al., 2018).

NSM_L/R_ and M3_L/R_ (NSM_L/R_ are *pezo-1*-expressing neurons), have been postulated to sense bacteria in the pharynx lumen, via their proprioceptive endings, and secrete serotonin in response to this mechanical stimulus (Avery, 1993, Avery and Thomas, 1997). Laser ablation of NSM_L/R_ in *unc-29* mutants leads to subtle changes in pharyngeal pumping rate; however, this was done while simultaneously ablating other pharyngeal motor neurons (M1, M2_L/R_, M3_L/R_, M5, and MI) (Avery, 1993). This approach could exert antagonistic effects on pumping rate, yielding a steady pharyngeal activity. Using the Screenchip™ system allowed us to reveal the potential roles of extrapharyngeal neurons expressing *pezo-1* (NSM_L/R_). Our results determined that proper function of PEZO-1 lengthened interpump intervals with 2 mM serotonin, in the absence of food. They further demonstrated that PEZO-1 modulated the feeding behavior of worms confronted with food of various mechanical properties (control and spaghetti-like bacteria). This led us to hypothesize that PEZO-1 is involved in food sensation and modulates pharyngeal pumping rate. Hence, like the mammalian ortholog PIEZO2, PEZO-1 is expressed in proprioceptive endings and is involved in stretch reflexes (Chesler et al., 2016, Woo et al., 2015). Nevertheless, it remains to be determined if mammalian PIEZO channels play a role in food sensation and/or the swallowing reflex.

Humans sense various organoleptic food qualities, such as visual aspects (color and shape), odorants through smell, and texture and flavor through tasting. In nematodes, there is a lack of understanding of what is sensed as food. Worms are able to filter particles from fluid in a size-dependent manner (Fang-Yen et al., 2009, Kiyama et al., 2012) and feeding is facilitated by attractive odors or suppressed by repellents (e.g., diacetyl, isoamyl alcohol, quinine) (Gruninger et al., 2008, Li et al., 2012). Others have demonstrated that worms prefer to feed from active (i.e., bacteria reproducing rapidly and emitting high levels of CO2), rather than inactive bacteria (Yu et al., 2015). We determined that *pezo-1* KO worms “choke” when presented with long and stiff spaghetti-like bacteria, whereas WT and GOF strains increase pharyngeal pumping when ingesting this elongated and rigid food. Therefore, we propose that the pharynx itself might be a sensory organ, as worms modify their pumping parameters when they sense solutions of different osmolarities or food with different textures and/or consistencies. We further hypothesize that worms can perceive changes in texture and adjust their pumping frequency through a mechanism requiring PEZO-1. Since *pezo-1* is not essential for *C. elegans* when cultured in standard laboratory conditions (e.g., monoaxenically on *E. coli* OP50), we wonder if in its natural biotic environment this mechanosensitive ion channel plays a crucial role, as it does in humans and *Drosophila*. Given that worms grow in microbe-rich and heterogenous environments (feeding from prokaryotes of the genera *Acetobacter*, *Gluconobacter*, and *Enterobacter*, for example) (Schulenburg and Felix, 2017), they might encounter bacteria of different dimensions and stiffness that would make *pezo-1* function more relevant to the worm’s ability to discriminate the food on which it grows best.

Why do *pezo-1* loss- and gain-of-function mutations cause similar behavioral phenotypes? Our data show that both *pezo-1* mutants (KO and GOF) increase the pumping frequency of the pharynx in different settings: serotonin exposure (albeit not statistically significant), high osmolarity, and ingestion of control bacteria. While it may seem counterintuitive at first, there are some scenarios in which too little or too much mechanosensation can be detrimental for animal behavior. In humans, *PIEZO2* LOF (premature stop codon) and GOF (missense mutation I802F) alleles cause joint contractures, skeletal abnormalities and alterations in muscle tone (Chesler et al., 2016, Coste et al., 2013, Yamaguchi et al., 2019). Only when feeding worms with spaghetti-like bacteria, were we able to uncover differences in the pharyngeal parameters between the LOF and the GOF mutants. Hence, we hypothesize that lacking the function of PEZO-1 significantly slows pharyngeal function when passing the lengthy and rigid bacteria from the pharynx to the gut.

Several requirements must be met for a channel to be considered mechanically-gated (Arnadottir and Chalfie, 2010). Accordingly, we found that *pezo-1* is expressed in the proprioceptive NSM neuron, knocking out *pezo-1* inhibits worm pharyngeal function when fed with elongated and stiff bacteria, engineering a single point mutation in the putative pore domain (R2373K) elicited similar inactivation, activation and deactivation delays that are reminiscent of the gating behavior reported for the human PIEZO1 R2456K (Bae et al., 2013), and expression of *pezo-1* (WT and GOF) confers mechanosensitivity to, otherwise naïve, Sf9 cells. We propose that PEZO-1 is a mechanosensitive ion channel given that the time it takes to reach half of the steady state currents ranges between 3.5 to 15 ms upon application of negative pressure. These are faster than activation times reported for the *Drosophila* phototransduction cascade, one of the most rapid second messenger cascades (Hardie, 2001). These combined efforts highlight the versatile functions of the PIEZO mechanosensitive channel family as well as the strength of the *C. elegans* model organism to reveal physiological functions.

Our findings revealing PEZO-1 as a mechanosensitive ion channel that modulates pharyngeal function raise several important questions. How does *pezo-1* modulate pumping behavior electrical activity? Does *pezo-1* equally enhance or inhibit the function of the pharyngeal hypodermal, gland and muscle cells, and neurons expressing this channel? Could *pezo-1* phenotypes be exacerbated if the gene function is nulled in a cell-specific manner? Does the slow deactivation and/or inactivation of the GOF mutant, determined at the patch-clamp level, account for the enhancement in pharyngeal function when worms are fed with bacteria? Does PEZO-1 require auxiliary subunits and/or the cytoskeleton for gating? Regardless of the answers, the plethora of physiological roles that this eukaryotic family of mechanosensitive ion channels play is outstanding. More experimental insight will be needed to grasp the full implications of *pezo-*1 in the physiology of *C. elegans*.

## Acknowledgements

The authors thank Dr. Julio F. Cordero-Morales, Dr. Andrés G. Vidal-Gadea, and Dr. Christopher E. Hopkins for critically reading the manuscript, as well as Dr. Rebeca Caires and Graduate Students Olufunke Falayi and Soumi Mazumdar for technical assistance. LX960 was provided by Dr. Kevin Collins (University of Miami). *C. elegans* (N2, DA572, and DA1051) and *E. coli* strains (OP50 and NA22) were obtained from the *Caenorhabditis* Genetics Center, which is funded by the NIH Office of Research Infrastructure Programs (P40 OD010440). This work was supported by the American Heart Association (16SDG26700010 to VV) and the National Institutes of Health (R01GM133845 to VV), as well as the Neuroscience Institute at UTHSC (Research Associate Matching Salary Support to JL).

## Author contributions

Conceptualization, VV; Formal Analysis, VV, JRMM, and LOR; Funding Acquisition, VV; Investigation, JRMM, LOR, JL, and BB; Methodology, VV and JRMM; Project Administration, VV; Supervision, VV; Writing - original draft, VV and JRMM; Writing - review & editing, VV.

## REFERENCES

Albertson, D. G. & Thomson, J. N. 1976. The pharynx of Caenorhabditis elegans. Philos Trans R Soc Lond B Biol Sci, 275, 299–325.

Albuisson, J., Murthy, S. E., Bandell, M., Coste, B., Louis-DIT-PICARD, H., Mathur, J., Feneant-THIBAULT, M., Tertian, G., De JAUREGUIBERRY, J. P., Syfuss, P. Y., Cahalan, S., Garcon, L., Toutain, F., Simon ROHRLICH, P., Delaunay, J., Picard, V., Jeunemaitre, X. & Patapoutian, A. 2013. Dehydrated hereditary stomatocytosis linked to gain-of-function mutations in mechanically activated PIEZO1 ion channels. Nat Commun, 4, 1884.

Alper, S. L. 2017. Genetic Diseases of PIEZO1 and PIEZO2 Dysfunction. Curr Top Membr, 79, 97–134.

Altun, Z. F. H., D.H. 2013. Epithelial system, interfacial cells. WormAtlas.

Arnadottir, J. & Chalfie, M. 2010. Eukaryotic mechanosensitive channels. Annu Rev Biophys, 39, 111–37.

Avery, L. 1993. Motor neuron M3 controls pharyngeal muscle relaxation timing in Caenorhabditis elegans. J Exp Biol, 175, 283–97.

Avery, L., Bargmann, C. I. & Horvitz, H. R. 1993. The Caenorhabditis elegans unc-31 gene affects multiple nervous system-controlled functions. Genetics, 134, 455–64.

Avery, L. & Horvitz, H. R. 1989. Pharyngeal pumping continues after laser killing of the pharyngeal nervous system of C. elegans. Neuron, 3, 473–85.

Avery, L. & Thomas, J. H. 1997. Feeding and Defecation. In: Nd, Riddle, D. L., Blumenthal, T., Meyer, B. J. & Priess, J. R. (eds.) C. elegans II. Cold Spring Harbor (NY).

Avery, L. & You, Y. J. 2012. C. elegans feeding. WormBook, 1–23.

Bae, C., Gnanasambandam, R., Nicolai, C., Sachs, F. & Gottlieb, P. A. 2013. Xerocytosis is caused by mutations that alter the kinetics of the mechanosensitive channel PIEZO1. Proc Natl Acad Sci U S A, 110, E1162–8.

Bai, X., Bouffard, J., Lord, A., Brugman, K., Sternberg, P. W., Cram, E. J. & Golden, A. 2020. Caenorhabditis elegans PIEZO channel coordinates multiple reproductive tissues to govern ovulation. Elife, 9.

Ben AROUS, J., Laffont, S. & Chatenay, D. 2009. Molecular and sensory basis of a food related two-state behavior in C. elegans. PLoS One, 4, e7584.

Brenner, S. 1974. The genetics of Caenorhabditis elegans. Genetics, 77, 71–94.

Caires, R., Bell, B., Lee, J., Romero, L. O., Vasquez, V. & Cordero-MORALES, J. F. 2021. Deficiency of Inositol Monophosphatase Activity Decreases Phosphoinositide Lipids and Enhances TRPV1 Function In Vivo. J Neurosci, 41, 408–423.

Caires, R., Sierra-VALDEZ, F. J., Millet, J. R. M., Herwig, J. D., Roan, E., Vasquez, V. & Cordero-MORALES, J. F. 2017. Omega-3 Fatty Acids Modulate TRPV4 Function through Plasma Membrane Remodeling. Cell Rep, 21, 246–258.

Chesler, A. T., Szczot, M., Bharucha-GOEBEL, D., Ceko, M., Donkervoort, S., Laubacher, C., Hayes, L. H., Alter, K., Zampieri, C., Stanley, C., Innes, A. M., Mah, J. K., Grosmann, C. M., Bradley, N., Nguyen, D., Foley, A. R., Le PICHON, C. E. & Bonnemann, C. G. 2016. The Role of PIEZO2 in Human Mechanosensation. N Engl J Med, 375, 1355–1364.

Coste, B. 2012. [Piezo proteins form a new class of mechanically activated ion channels]. Med Sci (Paris), 28, 1056–7.

Coste, B., Houge, G., Murray, M. F., Stitziel, N., Bandell, M., Giovanni, M. A., Philippakis, A., Hoischen, A., Riemer, G., Steen, U., Steen, V. M., Mathur, J., Cox, J., Lebo, M., Rehm, H., Weiss, S. T., Wood, J. N., Maas, R. L., Sunyaev, S. R. & Patapoutian, A. 2013. Gain-of-function mutations in the mechanically activated ion channel PIEZO2 cause a subtype of Distal Arthrogryposis. Proc Natl Acad Sci U S A, 110, 4667–72.

Coste, B., Mathur, J., Schmidt, M., Earley, T. J., Ranade, S., Petrus, M. J., Dubin, A. E. & Patapoutian, A. 2010. Piezo1 and Piezo2 are essential components of distinct mechanically activated cation channels. Science, 330, 55–60.

Cox, C. D., Bavi, N. & Martinac, B. 2019. Biophysical Principles of Ion-Channel-Mediated Mechanosensory Transduction. Cell Rep, 29, 1–12.

Dent, J. A., Davis, M. W. & Avery, L. 1997. avr-15 encodes a chloride channel subunit that mediates inhibitory glutamatergic neurotransmission and ivermectin sensitivity in Caenorhabditis elegans. EMBO J, 16, 5867–79.

Douguet, D. & Honore, E. 2019. Mammalian Mechanoelectrical Transduction: Structure and Function of Force-Gated Ion Channels. Cell, 179, 340–354.

Fang-YEN, C., Avery, L. & Samuel, A. D. 2009. Two size-selective mechanisms specifically trap bacteria-sized food particles in Caenorhabditis elegans. Proc Natl Acad Sci U S A, 106, 20093–6.

Ge, J., Li, W., Zhao, Q., Li, N., Chen, M., Zhi, P., Li, R., Gao, N., Xiao, B. & Yang, M. 2015. Architecture of the mammalian mechanosensitive Piezo1 channel. Nature, 527, 64–9.

Geffeney, S. L. & Goodman, M. B. 2012. How we feel: ion channel partnerships that detect mechanical inputs and give rise to touch and pain perception. Neuron, 74, 609–19.

Geron, M., Kumar, R., Zhou, W., Faraldo-GOMEZ, J. D., Vasquez, V. & Priel, A. 2018. TRPV1 pore turret dictates distinct DkTx and capsaicin gating. Proc Natl Acad Sci U S A, 115, E11837–E11846.

Gottlieb, P. A., Bae, C. & Sachs, F. 2012. Gating the mechanical channel Piezo1: a comparison between whole-cell and patch recording. Channels (Austin), 6, 282–9.

Gruninger, T. R., Gualberto, D. G. & Garcia, L. R. 2008. Sensory perception of food and insulin-like signals influence seizure susceptibility. PLoS Genet, 4, e1000117.

Guo, Y. R. & Mackinnon, R. 2017. Structure-based membrane dome mechanism for Piezo mechanosensitivity. Elife, 6.

Hamilton, E. S., Schlegel, A. M. & Haswell, E. S. 2015. United in diversity: mechanosensitive ion channels in plants. Annu Rev Plant Biol, 66, 113–37.

Hardie, R. C. 2001. Phototransduction in Drosophila melanogaster. J Exp Biol, 204, 3403–9.

Hart, A. C. 2006. WormBook.

Hasse, S., Hyman, A. A. & Sarov, M. 2016. TransgeneOmics--A transgenic platform for protein localization based function exploration. Methods, 96, 69–74.

Hoflich, J., Berninsone, P., Gobel, C., Gravato-NOBRE, M. J., Libby, B. J., Darby, C., Politz, S. M., Hodgkin, J., Hirschberg, C. B. & Baumeister, R. 2004. Loss of srf-3-encoded nucleotide sugar transporter activity in Caenorhabditis elegans alters surface antigenicity and prevents bacterial adherence. J Biol Chem, 279, 30440–8.

Hou, S., Jia, Z., Kryszczuk, K., Chen, D., Wang, L., Holyst, R. & Feng, X. 2020. Joint effect of surfactants and cephalexin on the formation of Escherichia coli filament. Ecotoxicol Environ Saf, 199, 110750.

Ikeda, R., Cha, M., Ling, J., Jia, Z., Coyle, D. & Gu, J. G. 2014. Merkel cells transduce and encode tactile stimuli to drive Abeta-afferent impulses. Cell, 157, 664–75.

Jang, H. & Bargmann, C. I. 2013. Acute behavioral responses to pheromones in C. elegans (adult behaviors: attraction, repulsion). Methods Mol Biol, 1068, 285–92.

Keane, J. & Avery, L. 2003. Mechanosensory inputs influence Caenorhabditis elegans pharyngeal activity via ivermectin sensitivity genes. Genetics, 164, 153–62.

Kim, S. E., Coste, B., Chadha, A., Cook, B. & Patapoutian, A. 2012. The role of Drosophila Piezo in mechanical nociception. Nature, 483, 209–12.

Kiyama, Y., Miyahara, K. & Ohshima, Y. 2012. Active uptake of artificial particles in the nematode Caenorhabditis elegans. J Exp Biol, 215, 1178–83.

Kung, C., Martinac, B. & Sukharev, S. 2010. Mechanosensitive channels in microbes. Annu Rev Microbiol, 64, 313–29.

Lee, K. S., Iwanir, S., Kopito, R. B., Scholz, M., Calarco, J. A., Biron, D. & Levine, E. 2017. Serotonin-dependent kinetics of feeding bursts underlie a graded response to food availability in C. elegans. Nat Commun, 8, 14221.

Lee, R. Y., Sawin, E. R., Chalfie, M., Horvitz, H. R. & Avery, L. 1999. EAT-4, a homolog of a mammalian sodium-dependent inorganic phosphate cotransporter, is necessary for glutamatergic neurotransmission in caenorhabditis elegans. J Neurosci, 19, 159–67.

Li, J., Hou, B., Tumova, S., Muraki, K., Bruns, A., Ludlow, M. J., Sedo, A., Hyman, A. J., Mckeown, L., Young, R. S., Yuldasheva, N. Y., Majeed, Y., Wilson, L. A., Rode, B., Bailey, M. A., Kim, H. R., Fu, Z., Carter, D. A., Bilton, J., Imrie, H., Ajuh, P., Dear, T. N., Cubbon, R. M., Kearney, M. T., Prasad, R. K., Evans, P. C., Ainscough, J. F. & Beech, D. J. 2014. Piezo1 integration of vascular architecture with physiological force. Nature, 515, 279–282.

Li, Z., Li, Y., Yi, Y., Huang, W., Yang, S., Niu, W., Zhang, L., Xu, Z., Qu, A., Wu, Z. & Xu, T. 2012. Dissecting a central flip-flop circuit that integrates contradictory sensory cues in C. elegans feeding regulation. Nat Commun, 3, 776.

Ma, S., Cahalan, S., Lamonte, G., Grubaugh, N. D., Zeng, W., Murthy, S. E., Paytas, E., Gamini, R., Lukacs, V., Whitwam, T., Loud, M., Lohia, R., Berry, L., Khan, S. M., Janse, C. J., Bandell, M., Schmedt, C., Wengelnik, K., Su, A. I., Honore, E., Winzeler, E. A., Andersen, K. G. & Patapoutian, A. 2018. Common PIEZO1 Allele in African Populations Causes RBC Dehydration and Attenuates Plasmodium Infection. Cell, 173, 443–455 e12.

Maksimovic, S., Nakatani, M., Baba, Y., Nelson, A. M., Marshall, K. L., Wellnitz, S. A., Firozi, P., Woo, S. H., Ranade, S., Patapoutian, A. & Lumpkin, E. A. 2014. Epidermal Merkel cells are mechanosensory cells that tune mammalian touch receptors. Nature, 509, 617–21.

Mango, S. E. 2007. The C. elegans pharynx: a model for organogenesis. WormBook, 1–26.

Martinac, B., Buechner, M., Delcour, A. H., Adler, J. & Kung, C. 1987. Pressure-sensitive ion channel in Escherichia coli. Proc Natl Acad Sci U S A, 84, 2297–301.

Mcmillin, M. J., Beck, A. E., Chong, J. X., Shively, K. M., Buckingham, K. J., Gildersleeve, H. I., Aracena, M. I., Aylsworth, A. S., Bitoun, P., Carey, J. C., Clericuzio, C. L., Crow, Y. J., Curry, C. J., Devriendt, K., Everman, D. B., Fryer, A., Gibson, K., Giovannucci UZIELLI, M. L., Graham, J. M., JR., Hall, J. G., Hecht, J. T., Heidenreich, R. A., Hurst, J. A., Irani, S., Krapels, I. P., Leroy, J. G., Mowat, D., Plant, G. T., Robertson, S. P., Schorry, E. K., Scott, R. H., Seaver, L. H., Sherr, E., Splitt, M., Stewart, H., Stumpel, C., Temel, S. G., Weaver, D. D., Whiteford, M., Williams, M. S., Tabor, H. K., Smith, J. D., Shendure, J., Nickerson, D. A., UNIVERSITY OF WASHINGTON CENTER FOR MENDELIAN, G. & Bamshad, M. J. 2014. Mutations in PIEZO2 cause Gordon syndrome, Marden-Walker syndrome, and distal arthrogryposis type 5. Am J Hum Genet, 94, 734–44.

Min, S., Oh, Y., Verma, P., Whitehead, S. C., Yapici, N., Van VACTOR, D., Suh, G. S. & Liberles, S. 2021. Control of feeding by Piezo-mediated gut mechanosensation in Drosophila. Elife, 10.

Murthy, S. E., Loud, M. C., Daou, I., Marshall, K. L., Schwaller, F., Kuhnemund, J., Francisco, A. G., Keenan, W. T., Dubin, A. E., Lewin, G. R. & Patapoutian, A. 2018. The mechanosensitive ion channel Piezo2 mediates sensitivity to mechanical pain in mice. Sci Transl Med, 10.

Niacaris, T. & Avery, L. 2003. Serotonin regulates repolarization of the C. elegans pharyngeal muscle. J Exp Biol, 206, 223–31.

Ohmachi, M., Sugimoto, A., Iino, Y. & Yamamoto, M. 1999. kel-1, a novel Kelch-related gene in Caenorhabditis elegans, is expressed in pharyngeal gland cells and is required for the feeding process. Genes Cells, 4, 325–37.

Pan, B., Akyuz, N., Liu, X. P., Asai, Y., Nist-LUND, C., Kurima, K., Derfler, B. H., Gyorgy, B., Limapichat, W., Walujkar, S., Wimalasena, L. N., Sotomayor, M., Corey, D. P. & Holt, J. R. 2018. TMC1 Forms the Pore of Mechanosensory Transduction Channels in Vertebrate Inner Ear Hair Cells. Neuron, 99, 736–753 e6.

Parpaite, T. & Coste, B. 2017. Piezo channels. Curr Biol, 27, R250–R252.

Pathak, M. M., Nourse, J. L., Tran, T., Hwe, J., Arulmoli, J., Le, D. T., Bernardis, E., Flanagan, L. A. & Tombola, F. 2014. Stretch-activated ion channel Piezo1 directs lineage choice in human neural stem cells. Proc Natl Acad Sci U S A, 111, 16148–53.

Raizen, D. M. & Avery, L. 1994. Electrical activity and behavior in the pharynx of Caenorhabditis elegans. Neuron, 12, 483–95.

Raizen, D. M., Lee, R. Y. & Avery, L. 1995. Interacting genes required for pharyngeal excitation by motor neuron MC in Caenorhabditis elegans. Genetics, 141, 1365–82.

Ranade, S. S., Woo, S. H., Dubin, A. E., Moshourab, R. A., Wetzel, C., Petrus, M., Mathur, J., Begay, V., Coste, B., Mainquist, J., Wilson, A. J., Francisco, A. G., Reddy, K., Qiu, Z., Wood, J. N., Lewin, G. R. & Patapoutian, A. 2014. Piezo2 is the major transducer of mechanical forces for touch sensation in mice. Nature, 516, 121–5.

Retailleau, K., Duprat, F., Arhatte, M., Ranade, S. S., Peyronnet, R., Martins, J. R., Jodar, M., Moro, C., Offermanns, S., Feng, Y., Demolombe, S., Patel, A. & Honore, E. 2015. Piezo1 in Smooth Muscle Cells Is Involved in Hypertension-Dependent Arterial Remodeling. Cell Rep, 13, 1161–1171.

Rode, B., Shi, J., Endesh, N., Drinkhill, M. J., Webster, P. J., Lotteau, S. J., Bailey, M. A., Yuldasheva, N. Y., Ludlow, M. J., Cubbon, R. M., Li, J., Futers, T. S., Morley, L., Gaunt, H. J., Marszalek, K., Viswambharan, H., Cuthbertson, K., Baxter, P. D., Foster, R., Sukumar, P., Weightman, A., Calaghan, S. C., Wheatcroft, S. B., Kearney, M. T. & Beech, D. J. 2017. Piezo1 channels sense whole body physical activity to reset cardiovascular homeostasis and enhance performance. Nat Commun, 8, 350.

Romero, L. O., Massey, A. E., Mata-DABOIN, A. D., Sierra-VALDEZ, F. J., Chauhan, S. C., Cordero-MORALES, J. F. & Vasquez, V. 2019. Dietary fatty acids fine-tune Piezo1 mechanical response. Nat Commun, 10, 1200.

Saotome, K., Murthy, S. E., Kefauver, J. M., Whitwam, T., Patapoutian, A. & Ward, A. B. 2018. Structure of the mechanically activated ion channel Piezo1. Nature, 554, 481–486.

Schindelin, J., Arganda-CARRERAS, I., Frise, E., Kaynig, V., Longair, M., Pietzsch, T., Preibisch, S., Rueden, C., Saalfeld, S., Schmid, B., Tinevez, J. Y., White, D. J., Hartenstein, V., Eliceiri, K., Tomancak, P. & Cardona, A. 2012. Fiji: an open-source platform for biological-image analysis. Nat Methods, 9, 676–82.

Schulenburg, H. & Felix, M. A. 2017. The Natural Biotic Environment of Caenorhabditis elegans. Genetics, 206, 55–86.

Shtonda, B. B. & Avery, L. 2006. Dietary choice behavior in Caenorhabditis elegans. J Exp Biol, 209, 89–102.

Singh, R. N. S., J.E. 1978. Some Observations On Moulting in Caenorhabditis Elegans. Nematologica, 24, 63–71.

Smit, R. B., Schnabel, R. & Gaudet, J. 2008. The HLH-6 transcription factor regulates C. elegans pharyngeal gland development and function. PLoS Genet, 4, e1000222.

Strange, K., Christensen, M. & Morrison, R. 2007. Primary culture of Caenorhabditis elegans developing embryo cells for electrophysiological, cell biological and molecular studies. Nat Protoc, 2, 1003–12.

Szczot, M., Liljencrantz, J., Ghitani, N., Barik, A., Lam, R., Thompson, J. H., Bharucha-GOEBEL, D., Saade, D., Necaise, A., Donkervoort, S., Foley, A. R., Gordon, T., Case, L., Bushnell, M. C., Bonnemann, C. G. & Chesler, A. T. 2018. PIEZO2 mediates injury-induced tactile pain in mice and humans. Sci Transl Med, 10.

Taylor, S. R., Santpere, G., Weinreb, A., Barrett, A., Reilly, M. B., Xu, C., Varol, E., Oikonomou, P., Glenwinkel, L., Mcwhirter, R., Poff, A., Basavaraju, M., Rafi, I., Yemini, E., Cook, S. J., Abrams, A., Vidal, B., Cros, C., Tavazoie, S., Sestan, N., Hammarlund, M., Hobert, O. & Miller, D. M., 3RD 2021. Molecular topography of an entire nervous system. Cell, 184, 4329–4347 e23.

Trojanowski, N. F., Raizen, D. M. & Fang-YEN, C. 2016. Pharyngeal pumping in Caenorhabditis elegans depends on tonic and phasic signaling from the nervous system. Sci Rep, 6, 22940.

Tsujimura, T., Ueha, R., Yoshihara, M., Takei, E., Nagoya, K., Shiraishi, N., Magara, J. & Inoue, M. 2019. Involvement of the epithelial sodium channel in initiation of mechanically evoked swallows in anaesthetized rats. J Physiol, 597, 2949–2963.

Van GILST, M. R., Hadjivassiliou, H., Jolly, A. & Yamamoto, K. R. 2005. Nuclear hormone receptor NHR-49 controls fat consumption and fatty acid composition in C. elegans. PLoS Biol, 3, e53.

Vidal-GADEA, A. G., Davis, S., Becker, L. & Pierce-SHIMOMURA, J. T. 2012. Coordination of behavioral hierarchies during environmental transitions in Caenorhabditis elegans. Worm, 1, 5–11.

Wang, P., Jia, Y., Liu, T., Jan, Y. N. & Zhang, W. 2020. Visceral Mechano-sensing Neurons Control Drosophila Feeding by Using Piezo as a Sensor. Neuron, 108, 640–650 e4.

Wang, S., Chennupati, R., Kaur, H., Iring, A., Wettschureck, N. & Offermanns, S. 2016. Endothelial cation channel PIEZO1 controls blood pressure by mediating flow-induced ATP release. J Clin Invest, 126, 4527–4536.

Woo, S. H., Lukacs, V., De NOOIJ, J. C., Zaytseva, D., Criddle, C. R., Francisco, A., Jessell, T. M., Wilkinson, K. A. & Patapoutian, A. 2015. Piezo2 is the principal mechanotransduction channel for proprioception. Nat Neurosci, 18, 1756–62.

Woo, S. H., Ranade, S., Weyer, A. D., Dubin, A. E., Baba, Y., Qiu, Z., Petrus, M., Miyamoto, T., Reddy, K., Lumpkin, E. A., Stucky, C. L. & Patapoutian, A. 2014. Piezo2 is required for Merkel-cell mechanotransduction. Nature, 509, 622–6.

Wu, J., Lewis, A. H. & Grandl, J. 2017a. Touch, Tension, and Transduction - The Function and Regulation of Piezo Ion Channels. Trends Biochem Sci, 42, 57–71.

Wu, J., Young, M., Lewis, A. H., Martfeld, A. N., Kalmeta, B. & Grandl, J. 2017b. Inactivation of Mechanically Activated Piezo1 Ion Channels Is Determined by the C-Terminal Extracellular Domain and the Inner Pore Helix. Cell Rep, 21, 2357–2366.

Yamaguchi, T., Takano, K., Inaba, Y., Morikawa, M., Motobayashi, M., Kawamura, R., Wakui, K., Nishi, E., Hirabayashi, S. I., Fukushima, Y., Kato, H., Takahashi, J. & Kosho, T. 2019. PIEZO2 deficiency is a recognizable arthrogryposis syndrome: A new case and literature review. Am J Med Genet A, 179, 948–957.

Yan, Z., Zhang, W., He, Y., Gorczyca, D., Xiang, Y., Cheng, L. E., Meltzer, S., Jan, L. Y. & Jan, Y. N. 2013. Drosophila NOMPC is a mechanotransduction channel subunit for gentle-touch sensation. Nature, 493, 221–5.

Yu, L., Yan, X., Ye, C., Zhao, H., Chen, X., Hu, F. & Li, H. 2015. Bacterial Respiration and Growth Rates Affect the Feeding Preferences, Brood Size and Lifespan of Caenorhabditis elegans. PLoS One, 10, e0134401.

Zarychanski, R., Schulz, V. P., Houston, B. L., Maksimova, Y., Houston, D. S., Smith, B., Rinehart, J. & Gallagher, P. G. 2012. Mutations in the mechanotransduction protein PIEZO1 are associated with hereditary xerocytosis. Blood, 120, 1908–15.

Zeng, W. Z., Marshall, K. L., Min, S., Daou, I., Chapleau, M. W., Abboud, F. M., Liberles, S. D. & Patapoutian, A. 2018. PIEZOs mediate neuronal sensing of blood pressure and the baroreceptor reflex. Science, 362, 464–467.

Zhao, Q., Zhou, H., Chi, S., Wang, Y., Wang, J., Geng, J., Wu, K., Liu, W., Zhang, T., Dong, M. Q., Wang, J., Li, X. & Xiao, B. 2018. Structure and mechanogating mechanism of the Piezo1 channel. Nature, 554, 487–492.

