## Supplementary Figures for "*C. elegans* PEZO-1 is a Mechanosensitive Ion Channel Involved in Food Sensation"

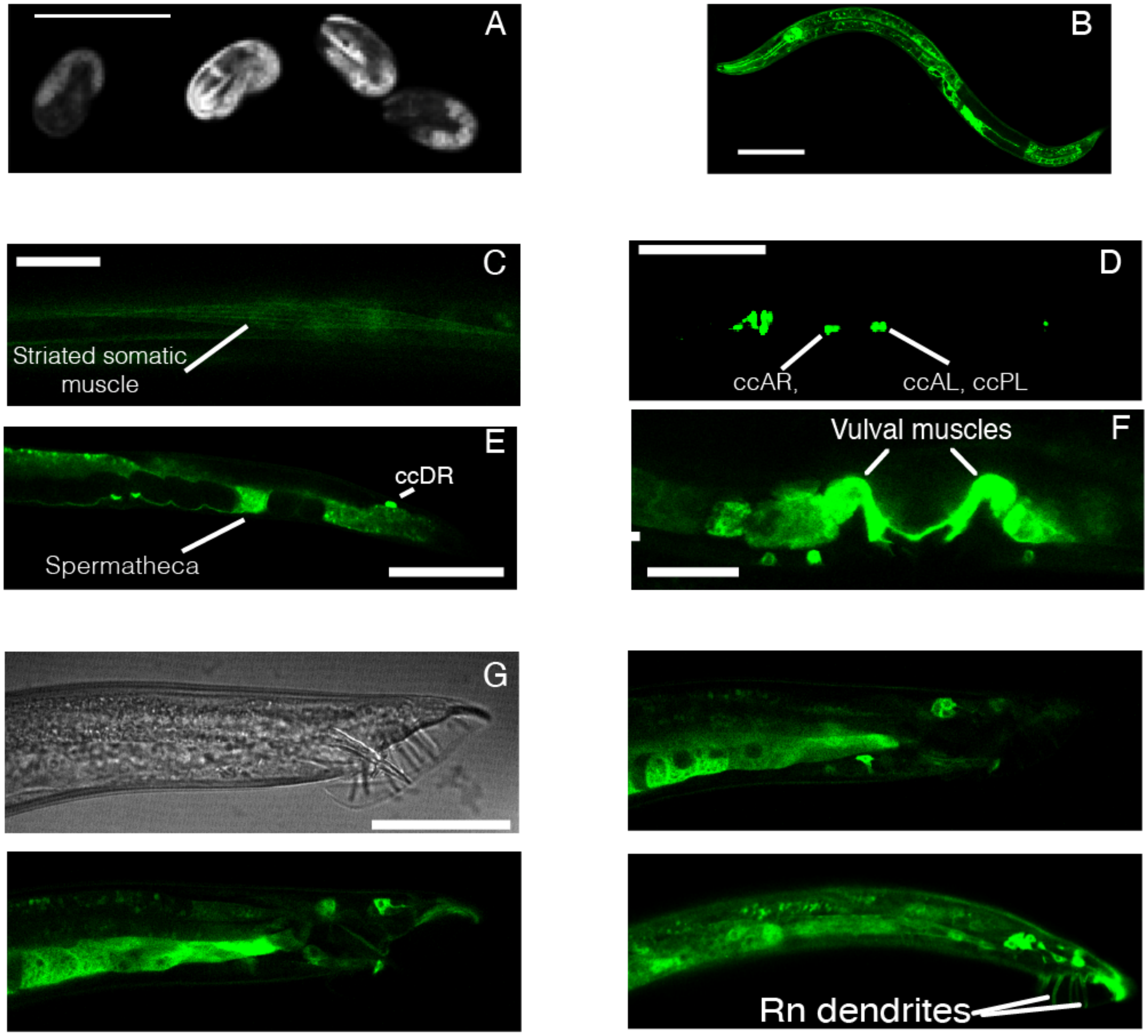


**Supplementary Figure 1**. ***pezo-1* is expressed in a variety of tissues in *C. elegans*.**
(**A**) Micrograph of *pezo-1*::*GFP* eggs. Scale bar represents 50 µm.
(**B**) Micrograph of an L4 *pezo-1*::GFP hermaphrodite expressing GFP. Scale bar represents 100 µm.
(**C**) Micrograph of a young adult *pezo-1*::*GFP* hermaphrodite expressing GFP in somatic muscles located in the anterior end of the animal. Scale bar represents 20 µm.
(**D**) Micrograph of a young adult *pezo-1*::*GFP* hermaphrodite expressing GFP in the coelomocytes (ccAR, ccAL, ccPL). Scale bar represents 100 µm.
(**E**) Micrograph of a larval *pezo-1*::*GFP* hermaphrodite expressing GFP in the spermatheca and in coelomocyte ccDR. Scale bar represents 100 µm.
(**F**) Micrograph of a young adult *pezo-1*::*GFP* hermaphrodite expressing GFP in the vulval muscle. Scale bar represents 20 µm.
(**G**) Micrographs (4) of a *pezo-1*::*GFP* adult male expressing GFP in the tail. Scale bar represents 50 µm. Rn: ray.

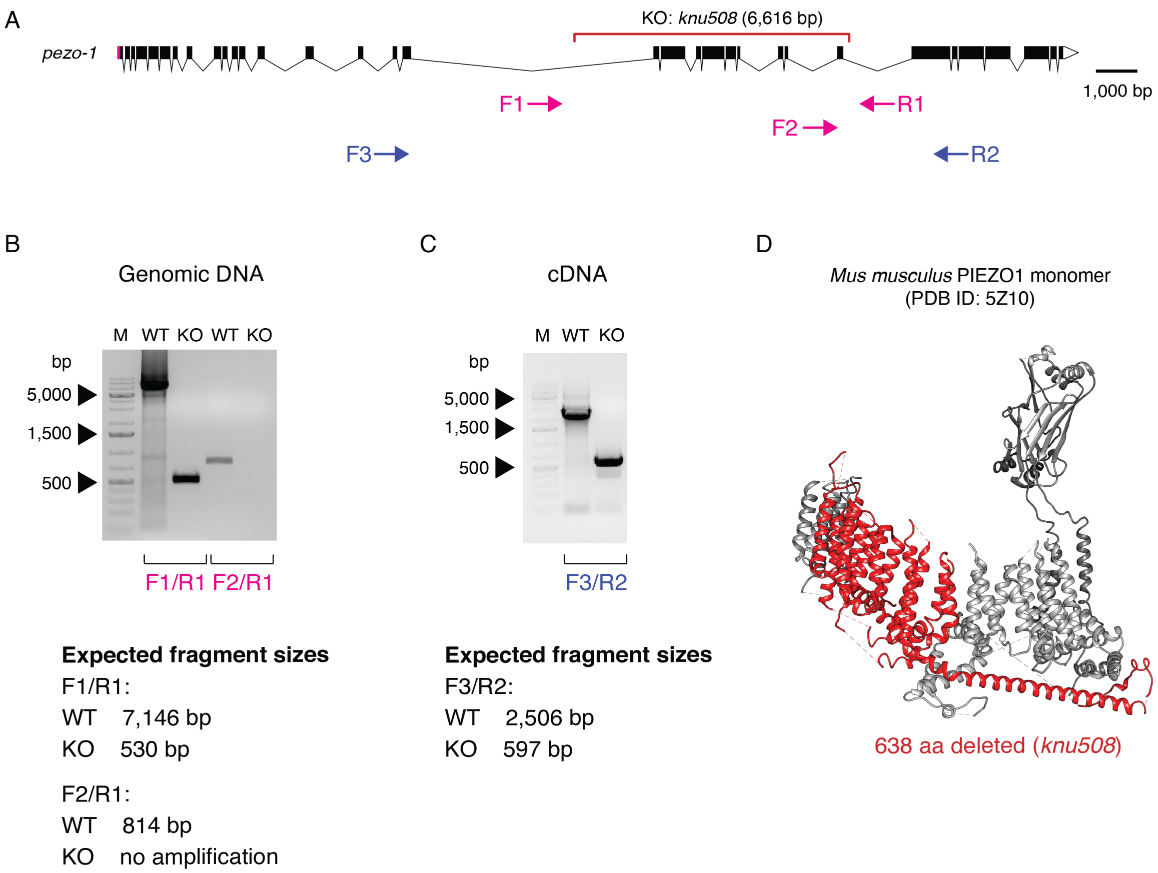


**Supplementary Figure 2. *pezo-1* KO validation.**

(**A**) *pezo-1* gene diagram according to wormbase.org v. WS280 made with Exon-Intron Graphic Maker (wormweb.org). Magenta rectangles and white triangles denote the 5' and 3' untranslated regions (UTR), respectively. Black rectangles denote exons and black lines denote introns. The red bracket denotes the *knu508* allele (a 6,616bp deletion) of the *pezo-1* KO strain. Magenta arrows labeled F1, F2, and R1 denote the positions of the oligos use for PCR amplification, and blue arrows F3 and R2 denote the positions of the oligos use for RT-PCR amplification.

**(B)** Agarose gel electrophoresis (1% agarose) of PCR amplified products using F1/R1 and F2/R1 PCR primer sets. Lane M, 1 kb Plus DNA (ThermoFisher Cat # SM1331/2) size marker. WT (N2) and KO (COP1553) refer to the worm strains used to extract genomic DNA.

**(C)** Agarose gel electrophoresis (1% agarose) of RT-PCR amplified products using F3/R2 primer sets. Lane M, 1 kb Plus DNA (ThermoFisher Cat # SM1331/2) size marker. WT (N2) and KO (COP1553) refer to the worm strains used to extract total RNA.

**(D)** Ribbon representation of *Mus musculus* PIEZO1 monomer (PDB ID: 5Z10; gray) highlighting the PEZO-1 corresponding residues (red) that were knocked using CRISPR to generate the *knu508* allele. PEZO-1 monomer ribbon diagram was made with UCSF Chimera v. 1.9.

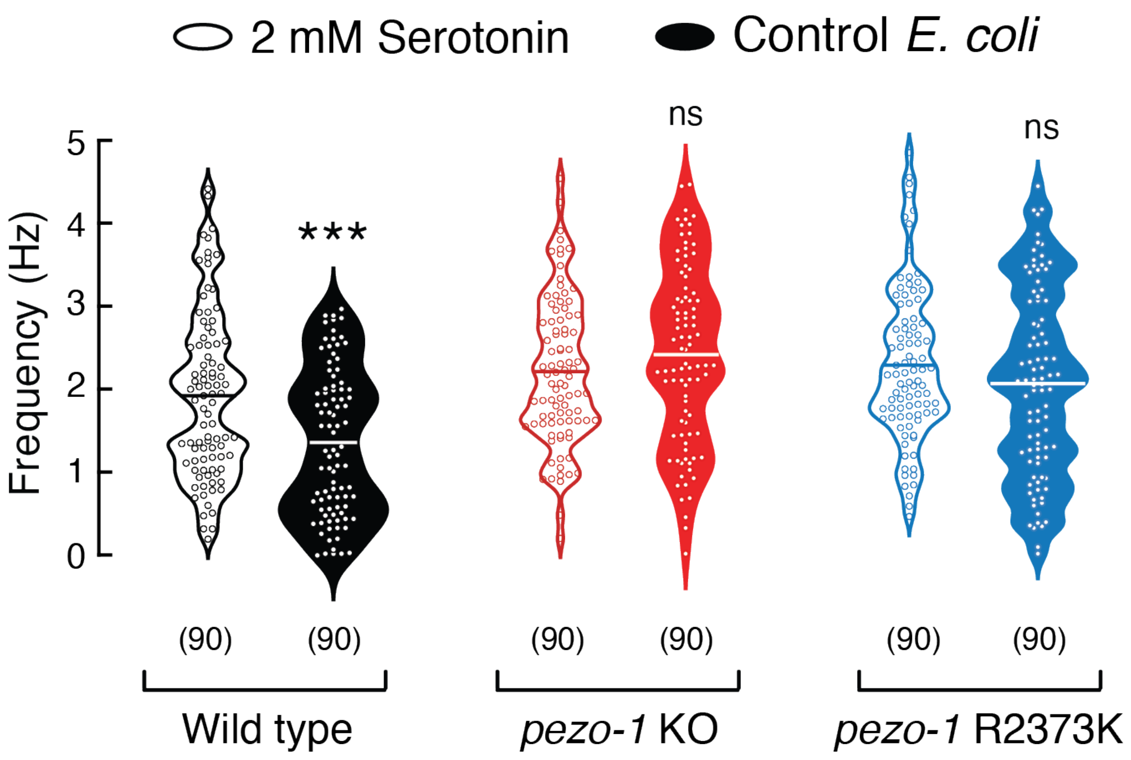


**Supplementary Figure 3. Comparison between *pezo-1* strains pumping frequencies elicited by serotonin or bacteria.**

Pharyngeal pumping frequencies depicted as violin plots with the means shown as horizontal bars, for WT (N2), *pezo-1* KO, and *pezo-1* R2373K strains at 2 mM serotonin concentration or when fed with control *E. coli*. n is denoted above the *x*-axis. Mann–Whitney test. Asterisks indicate values significantly different (***$p < 0.001$) and ns indicates not significantly different.

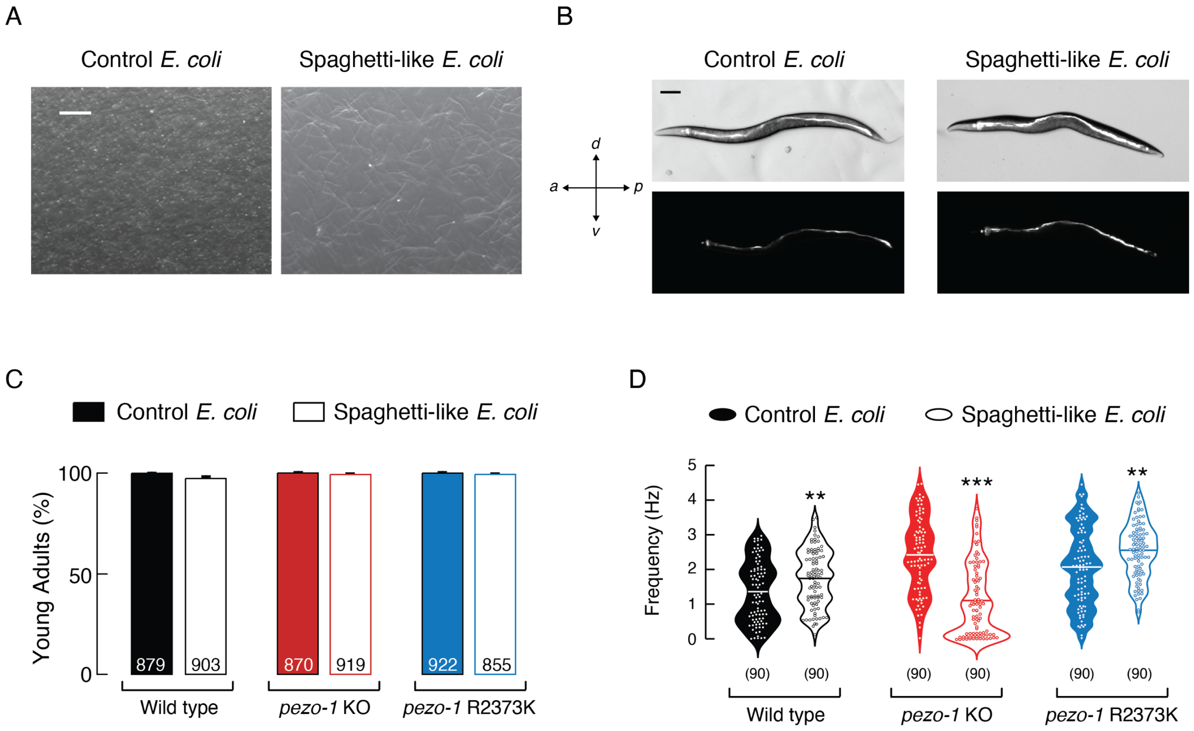


**Supplementary Figure 4. Worms eat spaghetti-like bacteria and reach adulthood.**
(**A**) Representative micrographs of control (left) and cephalexin-treated (right; spaghetti like) *Escherichia coli* cultures. Scale bar represents 200 µm.
**(B)** Representative micrographs of adult worms fed with control (left) and cephalexin-treated (right; spaghetti-like) *Escherichia coli* dyed with with 2 µM DiI. Scale bar represents 100 µm.
**(C)** WT (N2), *pezo-1* KO, and *pezo-1* R2373K adult proportion after three days of seeding eggs on NGM plates with control or spaghetti-like bacteria, as determined by worm images. Animals that reached adulthood were counted in each trial, and results were compared across four trials. n is denoted inside the bars.
(**D**) Pharyngeal pumping frequencies depicted as violin plots with the means shown as horizontal bars, for WT (N2), *pezo-1* KO, and *pezo-1* R2373K strains when fed with control or cephalexin-treated *E. coli* (spaghetti-like bacteria). n is denoted above the *x*-axis. Mann–Whitney test. Asterisks indicate values significantly different (***$p < 0.001$ and **$p < 0.01$).

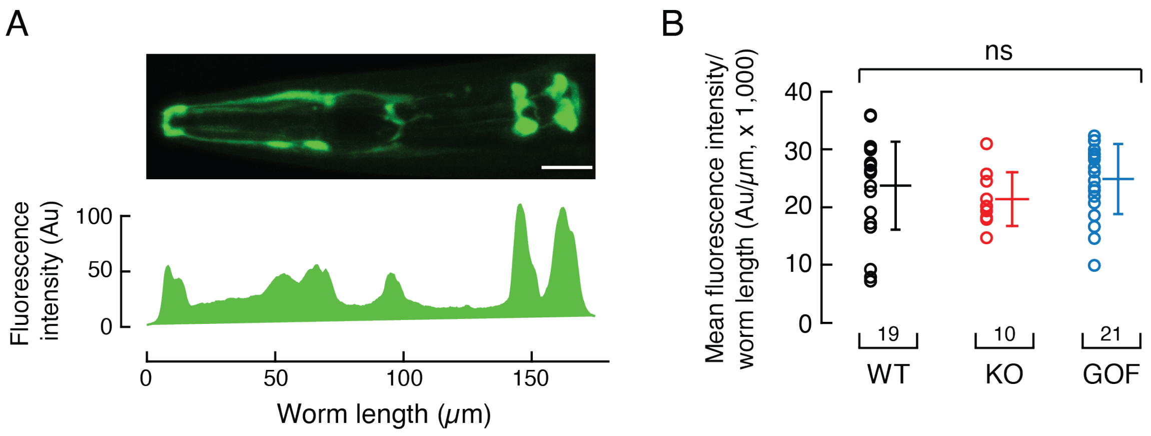


**Supplementary Figure 5. Fluorescence intensity of cells expressing *pezo-1::GFP* from WT, *pezo*-1 KO, and *pezo-1* R2373K strains.**

**(A)** Top, representative fluorescence micrograph of the anterior end of a young adult *pezo-1*::*GFP* hermaphrodite highlighting the GFP reporter expression in pharynx structures and gland cells. Scale bar represents 20 µm. Bottom, representative fluorescence intensity profile obtained from top micrograph.

**(B)** Mean/scatter-dot plot representing the mean fluorescence intensity per worm length of *pezo-1::GFP* from WT, *p*ezo-1 KO, and *pezo-1* R2373K strains. One-way ANOVA.n is denoted above the *x*-axis. ns indicates not significantly different.

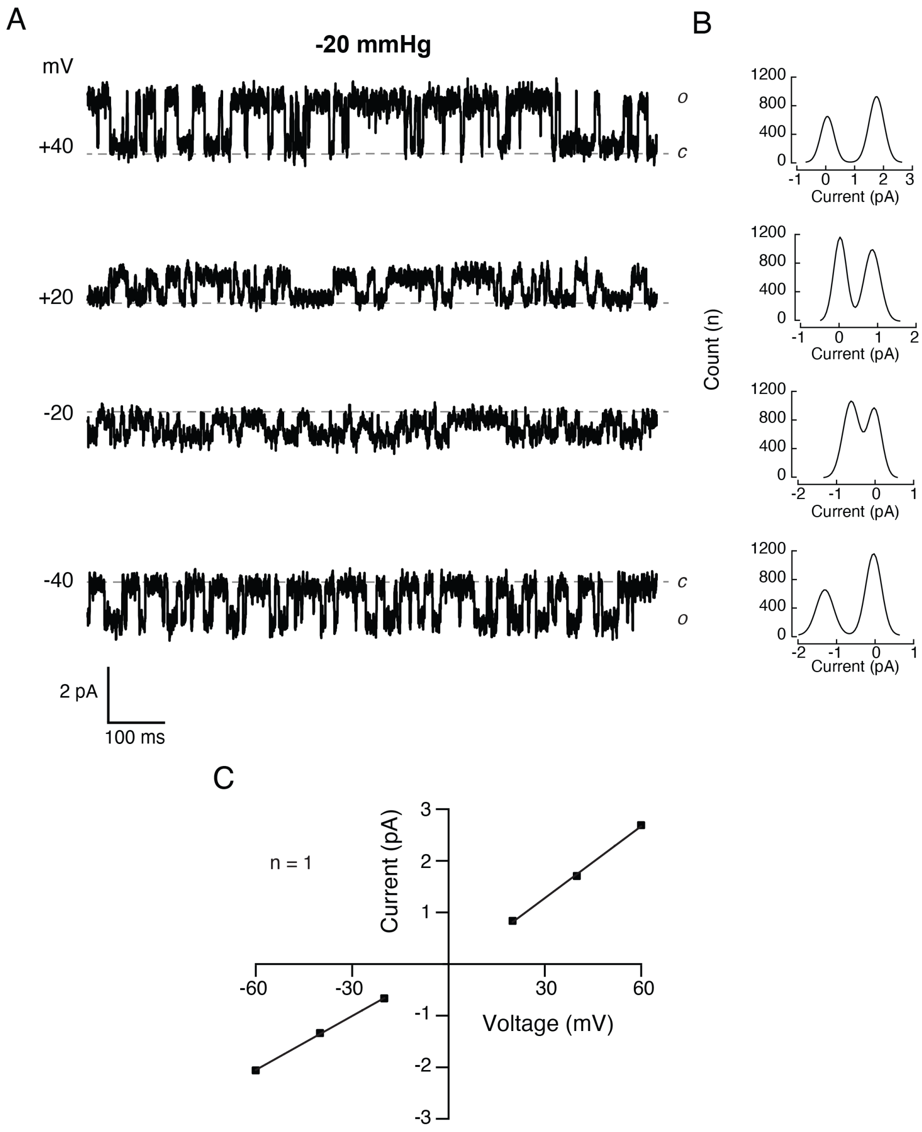


**Supplementary Figure 6. Mechanosensitive channel currents of cells expressing *pezo-1::GFP*.**
**(A)** Representative single-channel trace recordings of WT cells expressing *pezo-1::GFP* in the cell-attached configuration. Channel openings were elicited by -20 mmHg of negative pressure at constant voltages. Closed and open states are labeled *c* and *o*, respectively.
**(B)** All-point amplitude histograms of pressure-evoked single-channel currents from recordings shown in A.
**(C)** Current-voltage relationship at constant pressure (-20 mmHg). Outward slope conductance 46.37 pS and inward slope conductance 34.85 pS.

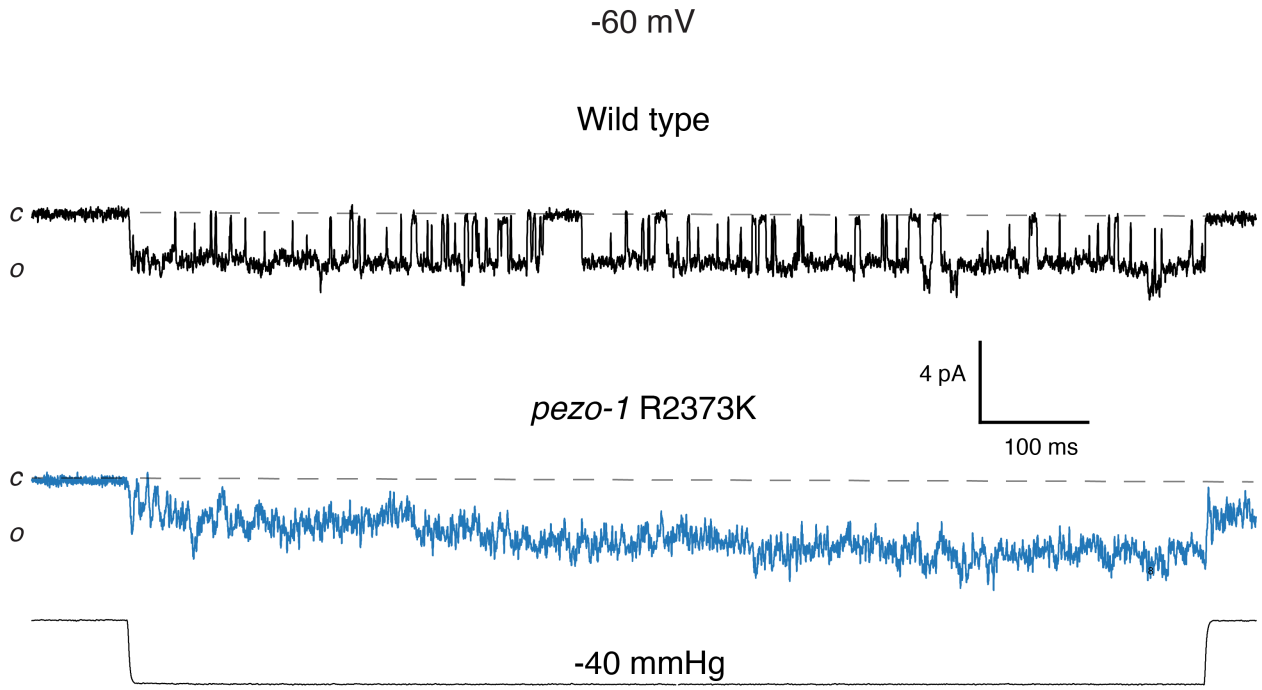


**Supplementary Figure 7**. Representative single-channel trace recordings of pressure-evoked currents from *pezo-1::GFP* cells expressing *pezo-1* WT and R2373K, in the cell-attached configuration. Channel openings were elicited by -40 mmHg at constant voltage -60 mV. These are the same records shown on Figure 7A-B used to generate the all-point amplitude histograms, filtered offline at 1 kHz to highlight single-channel events. Closed and open states are labeled *c* and *o*, respectively.

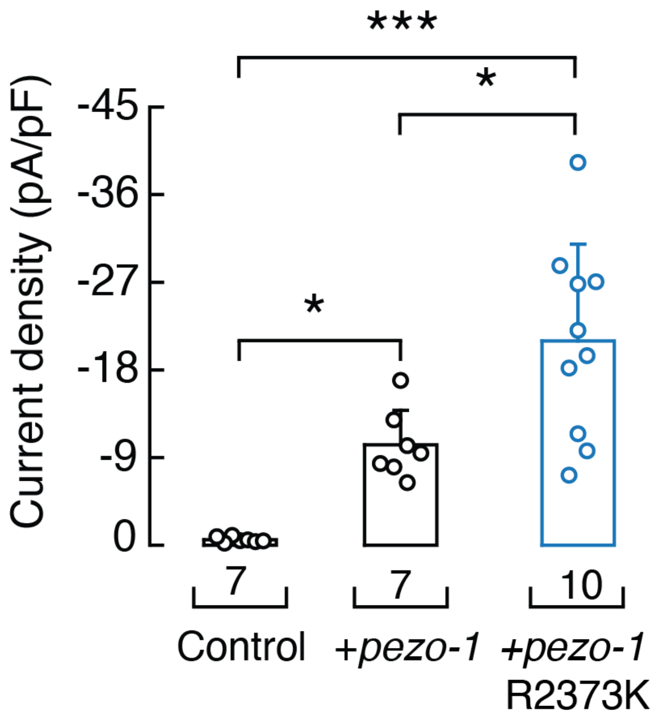


**Supplementary Figure 8**. Current densities elicited by maximum displacement of Sf9 cells control, expressing *pezo-1* WT or R2373K. n is denoted above the *x*-axis. One-way ANOVA and Tukey-Kramer multiple comparisons test. Asterisks indicate values significantly different (***$p < 0.001$ and *$p < 0.05$).

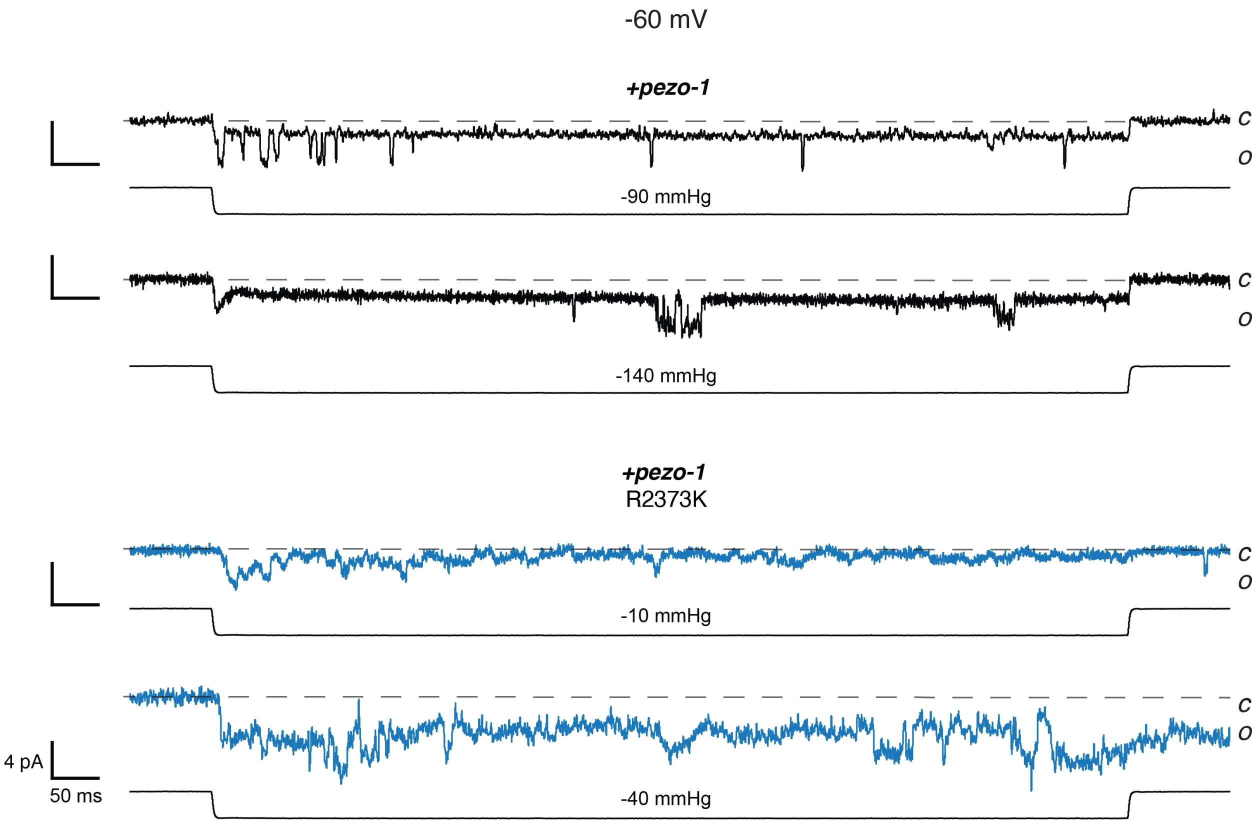


**Supplementary Figure 9**. Representative trace recordings of pressure-evoked currents from Sf9 cells expressing *pezo-1* WT or R2373K, recorded in the inside-out configuration. Channel openings were elicited by negative pressure at -60 mV. Traces were filtered offline at 1 kHz to highlight single-channel events. Closed and open states are labeled *c* and *o*, respectively.
